# High CD44 expression identifies rare chemoresistant leukemic cells endowed with enhanced E-Selectin binding in T-ALL

**DOI:** 10.1101/2023.12.20.572048

**Authors:** Julien Calvo, Irina Naguibneva, Anthony Kypraios, Benjamin Uzan, Baptiste Gaillard, Lea Bellenger L, Laurent Renou, Christophe Antoniewski, Helene Lapillone, Arnaud Petit, Paola Ballerini, Stéphane JC. Mancini, Tony Marchand, Jean-François Peyron, Françoise Pflumio

**Author notes:** I.N. and A.K. contributed equally to this study. J.C. and F.P. contributed equally to this study. Correspondence: Julien Calvo, Laboratoire des cellules Souches Hématopoïétiques et Leucémiques, UMR1274, IRCM, CEA, 18 route du panorama, 92260, Fontenay-aux-Roses, France. **Data sharing statement :** For original data, please contact.

## Abstract

T-cell acute lymphoblastic leukemia (T-ALL) is a hematopoietic malignancy characterized by an increased proliferation and incomplete maturation of T-cell progenitors. Despite therapeutic improvements, relapses are often of bad prognosis. Therapeutic vulnerabilities and chemoresistance mechanisms arising from cell plasticity induced by the bone marrow (BM) microenvironment remain an important field of investigation. Employing single cell RNA sequencing (scRNAseq) of human T-ALL cells recovered from adipocyte-rich and -poor BM, a distinct leukemic stem cell (LSC) population defined by quiescence and elevated CD44 level (Ki67^neg/low^CD44^high^) expression is identified in both territories. *In vivo* chemotherapy demonstrated that the LSC population evades drug treatment. Patient sample analyses confirmed the presence of Ki67^neg/low^CD44^high^ LSC both at diagnosis and relapse that displayed a specific transcriptomic signature. Interestingly, the intense expression of CD44 in T-ALL Ki67^neg/low^LSC was associated with E-selectin binding. Importantly, when 39 human T-ALL samples were analyzed, the E-selectin binding ability was found significantly higher in Relapse/Refractory compared to drug-sensitive patients. These findings characterize a T-ALL LSC population with chemoresistant properties and shade light on new strategies for prognostic stratification while opening avenues for novel therapeutic options.

## Introduction

T-cell Acute Lymphoblastic Leukemia (T-ALL) is characterized by the accumulation of genetic lesions that induce the differentiation arrest, survival and aberrant proliferation of immature T-cell progenitors^1,2^. Accounting for 15% of pediatric and 25% of adult cases of ALL, T-ALL has witnessed notable advancements in prognosis due to intensive chemotherapy. However, relapses still occur in 20% of pediatric and 50% of adult patients, often with a dismal outcome^3^. Identifying biomarkers indicative of relapse is imperative for precise patient classification based on risk of relapse. By taking advantage of these biomarkers, it is of paramount importance to uncover altenative vulnerabilities in resistant T-ALL cells in order to develop new selective drugs.

Leukemic cells infiltrate different tissues including the BM which provides an important microenvironment for T-ALL expansion. In there, leukemic cells take advantage of several BM-secreted factors described to support the initiation/spread of Acute Myeloid or Lymphoid Leukemia, such as CXCL12^4–7^, Interleukins 7 and 18, Insulin Growth Factor 1, or Notch-ligands^8^. Besides these soluble factors, multiple cells present within the BM niches such as hematopoietic cells, fibroblasts, osteoblasts/osteoclasts and neurons also interact with leukemic cells^9,10^. Adipocytes, which constitute Bone Marrow Adipose Tissue (BMAT), are rare in young bones but are massively increased by aging or radio/chemotherapy^11–14^. It has been demonstrated that adipocytes promote survival and/or proliferation of leukemic cells through 1) fatty acid assimilation^15–17^, 2) intense synthesis of L-asparagine and glutamine^18^ and 3) modulation of ROS production^19^. Due to these direct effects, adipocytes disturb the BM microenvironment. Indeed, adipocytes are negative regulators of the vasculature^11^, that help to generate hypoxic areas known to protect leukemic cells against chemotherapy^20,21^. Interactions between adipocytes and leukemic cells may thus be a key question in patient care. In fact, we and others previously showed that ALL cells infiltrate BMAT-rich sites in which they display decreased metabolism and translational processes, an accumulation in the G0 cell cycle phase, together with a chemoresistance phenotype^14,22^. This resistance cell state is plastic, since when BMAT-seeded leukemic cells are delocated in regular hematopoietic BM niches, they switch back to a proliferative/activated state^22^. These results indicate an attractive therapeutic window to overcome BMAT-induced chemoresistance.

Given the recognized role of the bone marrow microenvironment in T-ALL chemoresistance, here we fully characterize the chemoresistant quiescent T-ALL cells from BMAT sites. Using single-cell RNA sequencing analysis (scRNAseq), we uncovered a distinct minor cell population (called T-ALL LSC) that bear high chemoresistance potential and is in a quiescent state (Ki67^low/neg^). The transcriptomic signature of LSC population shares highly significant similarities with resting B-ALL cells previously described^23^. Furthermore, LSC population prominently expresses standard CD44 and increased presence of the tetrasaccharide carbohydrate termed sialylated Lewis X (sLeX) motif, allowing the interaction with E-selectin^24^. ScRNAseq analysis of primary T-ALL samples demonstrated the presence of Ki67^low/neg^CD44^high^ LSC population both at diagnosis and relapse, underscoring their physiological relevance. Furthermore, CD44/E-selectin binding was found enhanced in patients with poor prognosis, offering potential prognostic stratification.

## Results

### Detection of a quiescent T-ALL cell cluster in adipocyte-poor and -rich BM

To characterize the leukemic cell diversity in different BMAT-poor/rich sites^14,22^, we isolated cells from BMAT-poor BM (Femur, Thorax) and BMAT-rich BM (Tail Vertebrae) (Supplementary Figure 1a), then purified human leukemic huCD45^+^ for scRNAseq analysis. After quality filtering we obtained 30 544 cells (see materials and methods, Supplementary Figure 1b) displaying a median of 2 113 genes/cell and 4 978 transcript/cell (Figure 1a). Unsupervised clustering using the PCA (Principal Component Analysis) dimensional space allowed the visualization of 8 different clusters of leukemic cells pooled from the Femur, Thorax and Tail Vertebrae on UMAP (*Uniform Manifold Approximation and Projection)* (Figure 1b). The top 20 differentially expressed genes per cluster have been defined (Figure 1c). While all clusters exhibited an equitable distribution of cells across BM territories, Cluster 4 notably comprised 91.2% of cells from yellow/BMAT-rich sites (Figure 1d, Supplementary Figure 1c-d). Further analysis identified 383 and 569 genes respectively up- and down-regulated with log2 Fold Change > 0.25 and an adjusted p-value < 0.05 (Supplementary Table 4, Supplementary Figure 1e). Analysis of the cell cycle status in the scRNAseq dataset unveiled that cells from Cluster 0 and 4 were predominantly in the G1 phase, whereas cells from the other Clusters were in the S/G2M phases (Figure 1e). Low mRNA counts used to distinguish G0 from G1 phases as well as the absence of *MKI67* expression indicated that Cluster 4 contained cells mainly in G0 phase (Figure 1f-g, Supplementary Figure 1f). In order to know whether Cluster 4 was only defined by cell cycle progression genes, a new clustering analysis was performed by excluding the Principal Components (PCs) associated with cell cycle-related genes. Cluster 4 still stood aside from the other clusters, indicating that this cluster does not only rely on cell cycle genes (Supplementary Figure 1g). Accordingly, we observed by flow cytometry analysis, that leukemic cells from Femur and Thorax Vertebrae/BMAT-poor BM are activated and cycling cells, as compared to cells from Tail Vertebrae/BMAT-rich BM, which are mostly in a G0/quiescent state (Figure 1h-i). Collectively, these findings underscore significant leukemic cell diversity in both red and yellow BM sites. Our single cell transcriptomic analysis further indicates that most leukemic cells isolated from the BMAT-rich/yellow BM sites constitute a homogeneous cluster associated with quiescence. Interestingly, a low (8.8%), but significant, proportion of leukemic cells from BMAT-poor sites shared expression profiles with those from BMAT-rich/red sites (Supplementary Figure 1c).

**Figure 1:**
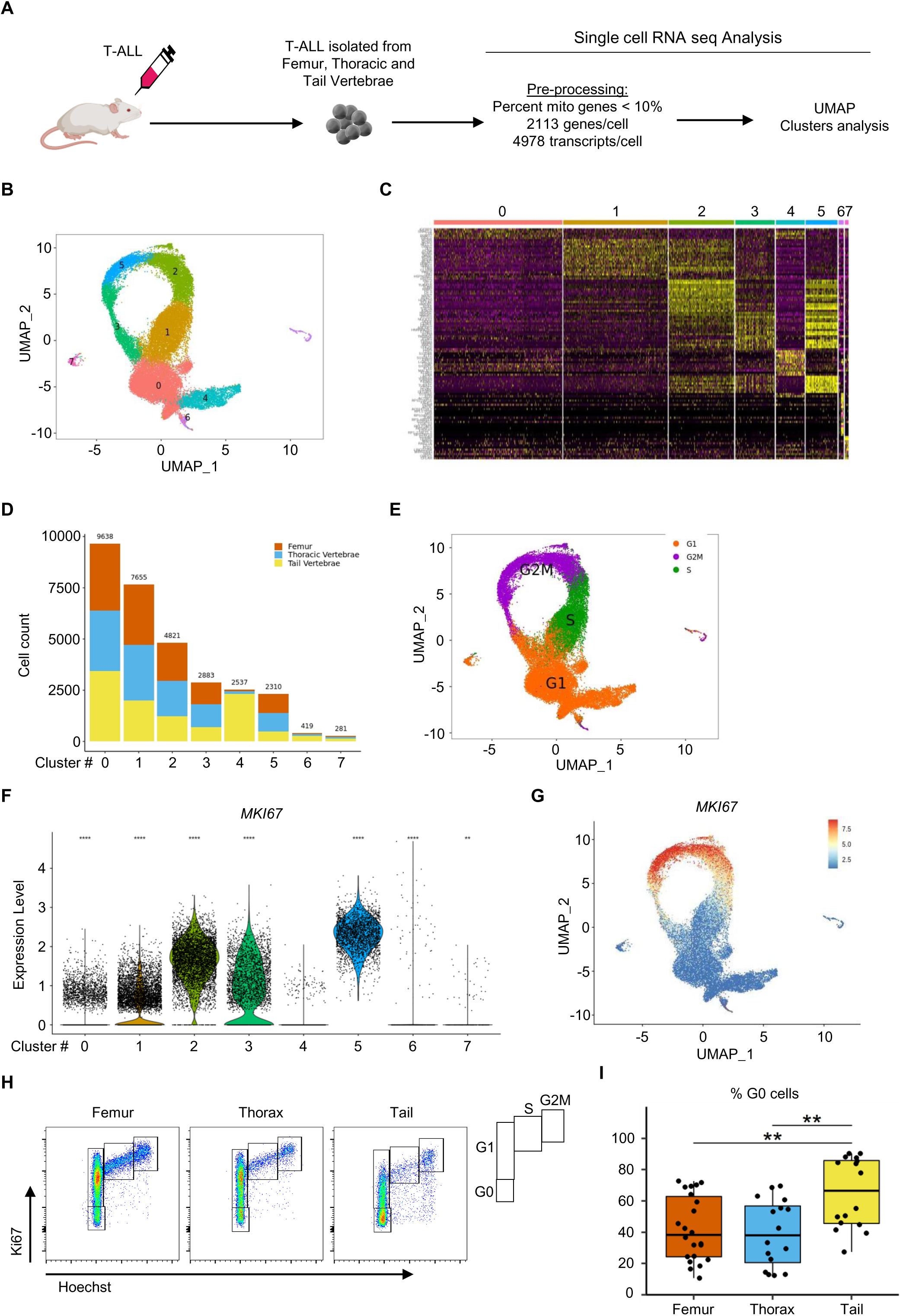
Characterization of leukemic cell diversity in adipocyte-rich/poor Bone Marrow. **A** Schematic overview of the study. **B** Uniform Manifold Approximation and Projection (UMAP) visualization of color-coded clustering of 30544 hT-ALL (M18) cells from Femur, Thoracic Vertebrae and Tail Vertebrae (4 mice). **C** Expression of 20 top differentially expressed genes (rows) across the cells (columns) in each cluster using DoHeatmap function. Colored bars correspond to the color-coded clustering (**B**). **D** Number of cells from Femur (orange), Thoracic Vertebrae (blue) and Tail Vertebrae (yellow) in each cluster. Data are presented in cumulative bar plots. **E** UMAP visualization of cell cycle progression using CellCycleScoring (detailled in Methods). **F-G** *MKI67* expression analysis for each cluster represented by Violin Plot associated with points denote values for each cell (**F**) and visualized on UMAP (**G**). Statistical significance was assessed by a Wilcoxon test for each cluster to the reference Cluster 4 (**** p < 0.0001; ** p < 0.01). **H-I** Cell cycle analysis by Flow Cytometry. Representative Ki67/Hoechst staining on humain CD45^+^/CD7^+^ T-ALL M106 cells from Femur, Thorax and Tail (**h**). Frequency of quiescent (G0) human CD45^+^/CD7^+^ T-ALL cells from femur (orange), thorax (blue) and tail vertebrae (yellow). Data are shown as box-and-whisker plots of the following numbers of mice for each group. Boxes indicate the 25th and 75th percentiles; whiskers display the range and horizontal lines in each bow represent the median. 4 PDX of hT-ALL tested from 16-22 mice (**i**). Statistical significance was assessed by Kruskal-Wallis test followed by Dunn’s multiple comparisons test (** p < 0.01). Human T-ALL samples are described in Supplementary Table 1.

### Quiescent T-ALL subpopulations are associated with high CD44 expression

The transcriptomic signature of Cluster 4 was further compared to the LRC (label-retaining cells) transcriptomic signature that previously defined quiescent B-ALL cells and is enriched in B-ALL MRD (Minimal Residual Disease) from patients^23^. We observed a significant enrichment of the LRC signature in cells from Cluster 4 (Figure 2a-b), underscoring a shared transcriptomic signature in quiescent and chemoresistant T/B-ALL. Over 18% of upregulated genes from the Cluster 4 signature are in common with upregulated genes from the LRC signature (Supplementary Figure 2a, Supplementary Table 5). In order to define the biological mechanisms that are associated with quiescent/resistant leukemic cells, we performed a Gene Set Enrichment Analysis (GSEA) of the upregulated genes from Cluster 4. Consistent with previous observations^14,22^, most of the basal cellular processes such as protein synthesis, mRNA catabolic processes and metabolism are downregulated in leukemic cells from Cluster 4 (Figure 2c and Supplementary Figure 2b-c, Supplementary Table 6a-c). The only significantly upregulated Gene Ontology (GO) biological processes highlighted in these resting leukemic cells were Cell Adhesion and Biological Adhesion processes whose Normalized Enrichment Score (NES) were 2.54 and 2.50 respectively (Figure 2c). Accordingly, Cellular Component analysis revealed that upregulated genes encode proteins related to “Localized to the cell membrane” GO term (Supplementary Figure 2b). We analyzed the mRNA and protein expression of the chemokine receptor CXCR4 given its pivotal role in T-ALL migration/homing and niche adhesion^6,7^. We observed that its expression was not higher in cells from the adipocyte-rich BM (Supplementary Figure 3). The top 5 genes of the Cell Adhesion mechanism enriched are S100A10, LGALS1, PTPRC, RIPOR2 and CD44 (Figure 2d, Supplementary Figure 4a-e). CD44 is an ubiquitous cell surface glycoprotein implicated in solid cancer and leukemia migration/adhesion^25–28^. Several CD44 variants, generated by alternative splicing, are known to be involved in cancer metastasis^29^. We determined which CD44 isoform is expressed in T-ALL cells using specific primers that target the different variants of *CD44* exons. We observed, by testing 12 Patient Derived Xenografts (PDX) obtained from 4 hT-ALL, that leukemic cells carry the *CD44* standard isoform (Supplementary Figure 5a) expressed at higher levels by cells from BMAT-rich/Tail Vertebrae compared to BMAT-poor BM (Figure 2e). Besides, surface CD44 protein was also higher on leukemic cells from BMAT-rich/Tail Vertebrae (Figure 2f), confirmed by Western Blot Analysis (Supplementary Figure 5b). We also observed that protein and mRNA levels were significantly correlated (r = 0.75 with p-value = 1.509 x 10^-14^, Supplementary Figure 5c) indicating that CD44 mRNA expression can be used as a reliable surrogate for protein levels in our T-ALL cell conditions.

**Figure 2:**
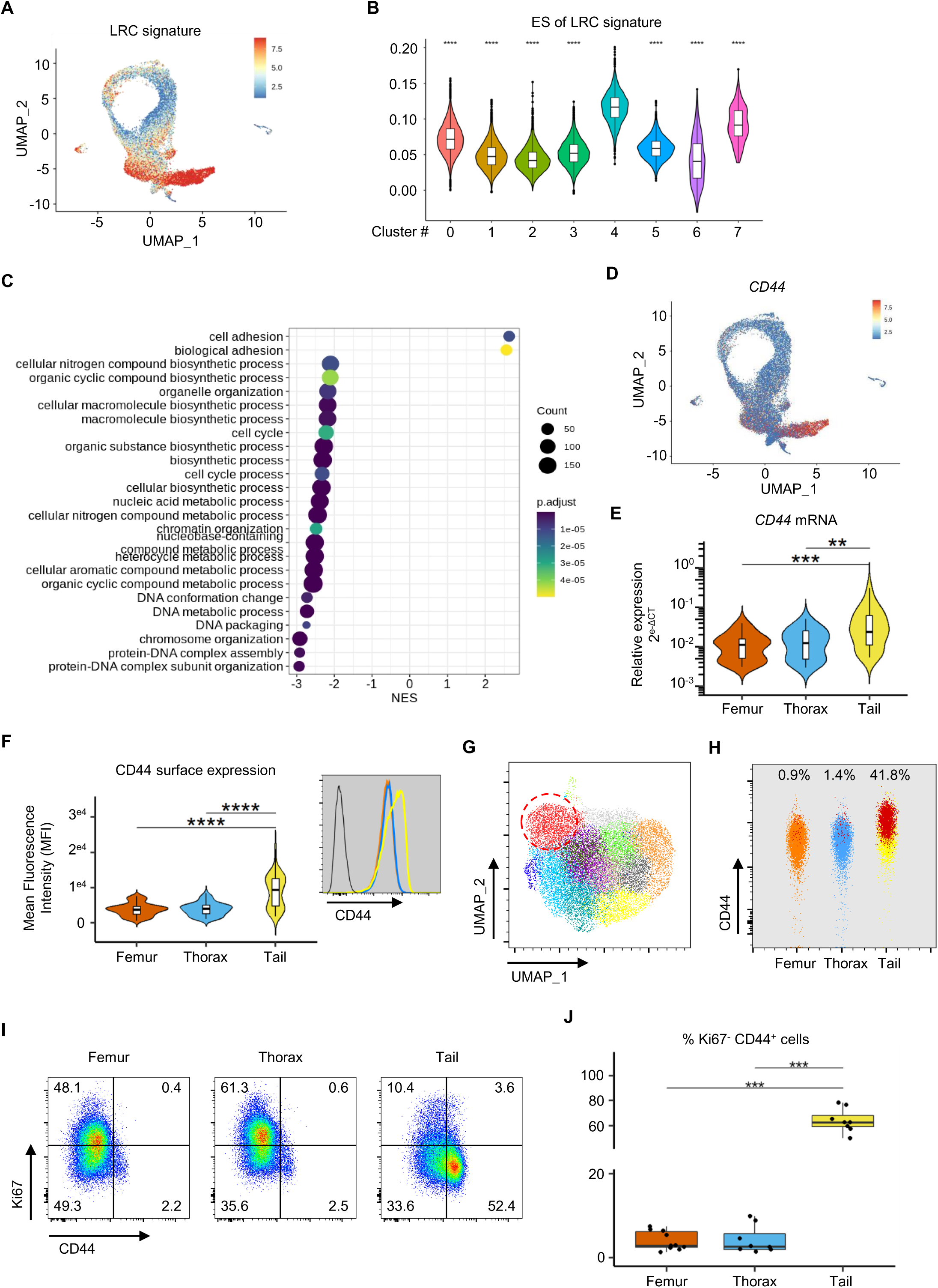
High expression of CD44 defines quiescent/dormant T-ALL cells. **A-B** Enrichment analysis (detailed in Methods) of LRC signature published by Ebinger et al on T-ALL cells from adipocyte-rich/poor BM visualized on UMAP (**A**) and represented for each cluster (**B**). **C** Gene Set Enrichment Analysis (GSEA) performed with significant upregulated genes from Cluster 4 of « Biological Process » by Gene Ontology annotation. Data shown as a Normalized Enrichment Score (NES) and significant processes are defined by FDR < 0.05 (color show the p.adjust level). **D** *CD44* expression analysis visualized on UMAP. **E** *CD44* relative expression analysis performed by RT-qPCR with T-ALL cells from Femur (orange), Thoracic Vertebrae (Blue) and Tail Vertebrae (Yellow). PDX of 15 hT-ALL samples (37 mice). **F** Mean Fluorescence Intensity (MFI) of CD44 expression performed by Flow Cytometry on human CD45^+^/CD7^+^ T-ALL cells from Femur (orange), Thorax (Blue) and Tail Vertebrae (Yellow). PDX of 19 hT-ALL samples (67 mice). Flow Cytometry analysis example obtained with M18-PDX, T-ALL cells unstained (black line), from Femur (orange line), Thorax (blue line) and Tail (yellow line) (inset). **E** and **F** Data are shown as violin and box-and-whisker plots. Violin indicate the density and boxes indicate the 25^th^ and 75^th^ percentiles; whiskers display the range and horizontal lines in each bow represent the median. **G** UMAP visualization of 21 600 hT-ALL (M18) cells from Femur, Thorax and Tail Vertebrae color-coded clustering based on FSC, SSC, CD7, CD45, CD4, CD8, CXCR4, CD44 and CD34 surface expression analyzed by Flow Cytometry. Red dashed line surrounds « Red cluster » which displays the highest CD44 expression. **H** Frequency of the « Red cluster » in each territory. **I-J** Flow Cytometry analysis of CD44 and Ki67 expression. Representative CD44/Ki67 staining of human CD45^+^/CD7^+^ T-ALL M106 cells from Femur, Thorax and Tail (**I**). Frequency of Ki67^neg/low^ CD44^high^ T-ALL cells from each territory. Data are shown as box-and-whisker plots of the following numbers of mice for each group. Boxes indicate the 25^th^ and 75^th^ percentiles; whiskers display the range and horizontal lines in each bow represent the median. 3 PDX (8-10 mice) (**J**). **E, F, J** Statistical significance was assessed by Kruskal-Wallis test followed by Dunn’s multiple comparisons test (** p < 0.01; *** p < 0.005). Human T-ALL samples are described in Supplementary Table 1.

### High CD44 expression is not related to NOTCH1 activation and TAL1 levels and is present in B-ALL

Activation of NOTCH1 signaling, a crucial regulator of T-ALL development/expansion^30^, drives the expression of critical genes such as *Myc*, *Hes1*, *pTalpha*, *Deltex1* and *CD44*^31,32^. To investigate whether high CD44 expression in cells from BMAT-rich site was due to NOTCH1 activation, we analyzed mRNA expression of several NOTCH1 target genes using scRNAseq data and RT-qPCR^30^. Data revealed no change of NOTCH1 pathway activation in leukemic cells retrieved from BMAT-poor compared to BMAT-rich sites (Supplementary Figure 6a-b). TAL1 transcription factor is recognized as a repressor of CD44 expression in CML (Chronic Myeloid Leukemia) cells^28^. From our results, we observed that CD44 expression is high in the Tail Vertebrae of mice, even though they are transplanted with TAL1 positive T-ALL samples (Supplementary Figure 6c-e). Analysis of TAL1 mRNA expression showed that its expression was similar whatever the analyzed BM territories, implying that TAL1 expression did not impact on CD44 expression in our conditions (Supplementary Figure 6f). In order to explore whether the observed high CD44 expression in BMAT-rich/yellow site was exclusive to T-ALL or could be extended to B-ALL, we analyzed CD44 expression in three B-ALL samples from different oncogenic groups (Supplementary Table 1). In line with the recent findings that B-ALL may recapitulate T-ALL phenotype in yellow BM^14^, we observed that CD44 expression, analyzed by RT-qPCR and flow cytometry, was also high in yellow BM compared to BMAT-poor/red sites (Supplementary Figure 7a-b). These last findings establish a consistent association of the CD44^high^ state with BMAT-rich/yellow BM for both T- and B-ALL cells.

### CD44^high^ leukemic cells are present in all BM sites, independently of BMAT richness

In parallel to scRNAseq, we also performed flow cytometry to further characterize leukemic cells. An unbiased UMAP clustering integrating 9 parameters including CD44 by multiparametric flow cytometry allowed to discriminate 12 clusters. Among these, a “Red-colored” Cluster (Figure 2g) consisted mainly of cells from Tail Vertebrae (95%, Supplementary Figure 8a) and accordingly displayed the highest CD44 levels (Supplementary Figure 8b). This CD44^high^ cell cluster (depicted as red dots in Figure 2h) represented 0.9%, 1.4% and 41.8% of the leukemic cells isolated from Femur, Thoracic and Tail Vertebrae respectively. These data highlight that BMAT-poor BM also harbors a minor leukemic cell population too, that matches to the major one recovered from BMAT-rich BM.

As with CD44^high^ leukemic cells from BMAT-rich sites (Figure 1h-i), we observed that CD44^high^ leukemic cells found in all BM territories are quiescent (Ki67^neg^) (Figure 2i-j). These findings are perfectly in line with the proportion of leukemic cells from BMAT-poor BM found in Cluster 4 (Supplementary Figure 1c). Taken together, our findings strongly suggest that high CD44 expression is associated with leukemic quiescence.

### *In vivo* chemotherapy efficacy is markedly attenuated whithin BMAT-rich compared to in BMAT-poor BM and CD44^high^ T-ALL cells exhibit inherent *in vivo* chemoresistant properties

To investigate whether the quiescent CD44^high^ T-ALL cells are endowed with chemoresistance, we implemented *in vivo* chemotherapy in NSG mice transplanted with human T-ALL cells^33–36^. Mice with substantial leukemia infiltration were treated with either vehicle or combination of Vincristine, Aracytidine, Dexamethasone and L-Asparaginase (VADA, Figure 3a and Supplementary Figure 9a-b). One week of treatment induced a drastic reduction of tumor burden in the spleen and peripheral blood (Supplementary Figure 9c-d). While the impact of VADA was significant across all BM sites, a clear trend to higher prounonced reduction in T-ALL cell burden within BMAT-poor rather than BMAT-rich areas emerged (Figure 3b and Supplementary Figure 9e). This observation highlights an uneven efficiency of drug treatment among BM sites, as previously observed *in vitro*^14,22^.

**Figure 3:**
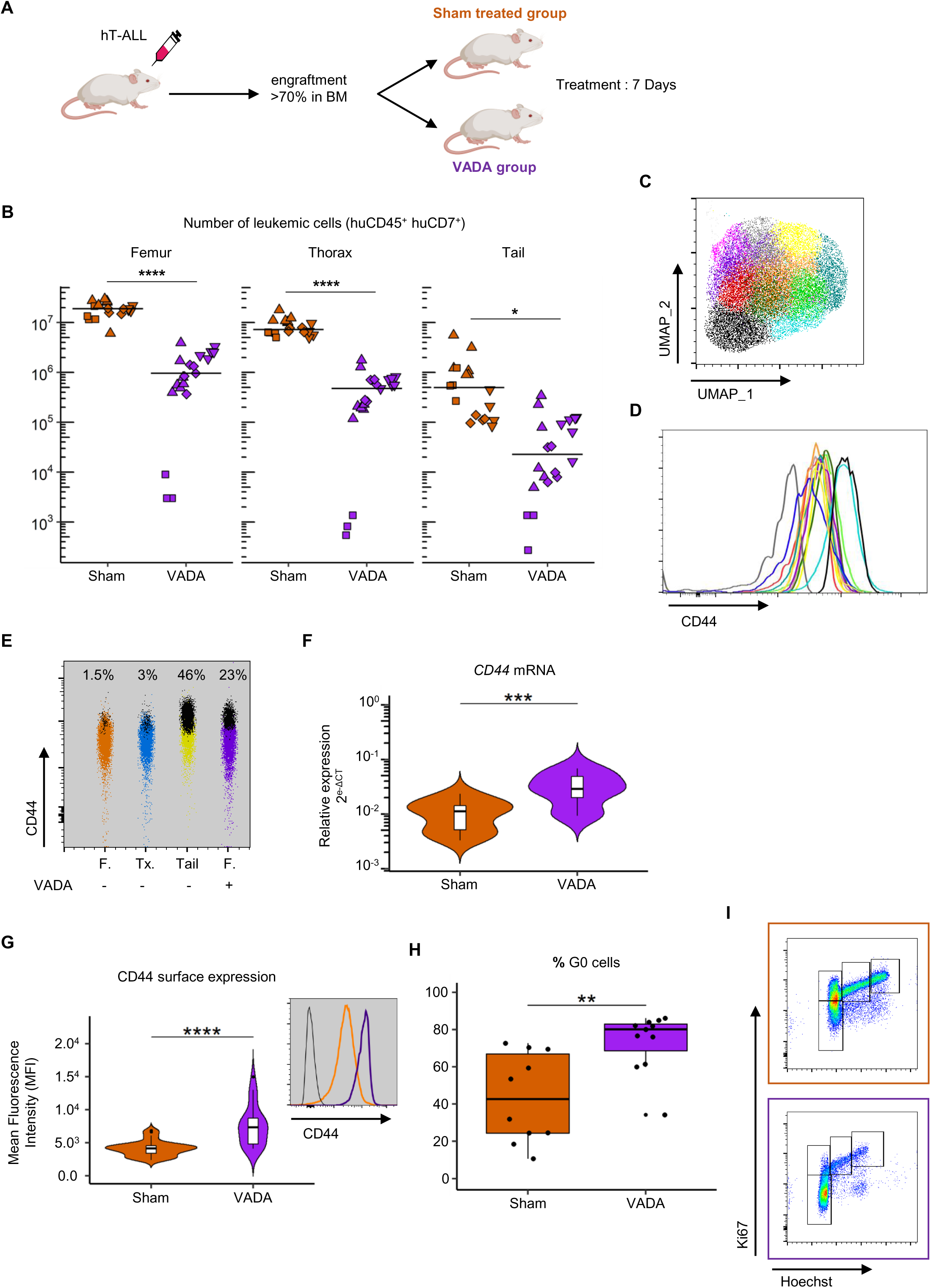
*In vivo* chemotherapy model to characterize Minimal Residual Disease Characterization of Minimal Residual Disease (MRD) from Femur after *in vivo* chemotherapy. A Schematic overview of the *in vivo* chemotherapy model. **B** Number of human CD45^+^/CD7^+^ T-ALL cells in each territory after Sham or VADA treatment. PDX of 4 hT-ALL samples (18 Sham and 21 VADA mice). Square, rhombus, triangle and inverted triangle symbols represent respectively M103, M69, M106 and M18 samples. **C-D** Cell heterogeneity evaluated by Flow Cytometry. UMAP visualization of 21 600 hT-ALL (M18) cells equally derived from Femur, Thorax and Tail Vertebrae of Sham mice mixed with 7 200 cells from Femur of VADA mice, using Phenograph color-coded clustering based on FSC, SSC, CD7, CD45, CD4, CD8, CXCR4, CD44 and CD34 surface expression (**C**). MFI of CD44 expression on the hT-ALL (M18) from each cluster (**D**). **E** Frequency of the « Black & Blue clusters » represented by black dots in Femur (orange), Thorax (blue), Tail (yellow) of Sham mice and Femur of VADA mice (purple). F *CD44* mRNA relative expression analyzed by RT-qPCR with purified hCD45^+^ hT-ALL cells from Femur from Sham (orange) and VADA mice (purple). PDX of 4 hT-ALL samples (19 Sham and 12 VADA mice). **G** MFI of CD44 expression analyzed by Flow Cytometry on human CD45^+^/CD7^+^ T-ALL cells from Femur from Sham (orange) and VADA mice (purple). PDX from 4 hT-ALL samples (35 Sham and 23 VADA mice). Flow Cytometry analysis example obtained with M106-PDX, T-ALL cells unstained (black line), from Femur Sham (orange line) and VADA mice (purple line) (inset). **F** and **G** Data are shown as violin and box-and-whisker plots. Violins indicate the density and boxes indicate the 25^th^ and 75^th^ percentiles; whiskers display the range and horizontal lines in each bow represent the median. **H-I** Quiescent analysis. Frequency of quiescent human CD45^+^/CD7^+^ T-ALL cells from Femur from Sham (orange) and VADA mice (purple). Data are shown as box-and-whisker plots of the following numbers of mice for each group. Boxes indicate the 25^th^ and 75^th^ percentiles; whiskers display the range and horizontal lines in each bow represent the median (**H**). PDX from 2 hT-ALL samples (10 Sham and 12 VADA mice). **i** Representative Ki67/Hoechst staining on human CD45^+^/CD7^+^ T-ALL (M69) cells from Femur from Sham (orange) and VADA mice (purple). **D, E, F** Statistical significance was assessed by a Mann & Withney test (** p < 0.01; *** p < 0.005; **** p < 0.001). Human T-ALL samples are described in Supplementary Table 1.

We next characterized residual BM T-ALL cells surviving VADA chemotherapy using comparative multi-parametric flow cytometry analysis of leukemic cells from Femur, Thoracic, Tail Vertebrae of Sham treated mice. An unbiased clustering approach using the UMAP algorithm coupled with Phenograph analysis revealed 12 clusters (Figure 3c-d), in which CD44^high^ leukemic cells are predominantly localized in two clusters (“Black&Blue”Clusters, Figure 3d). As shown in Figure 2h for untreated PDX, the proportion of the “Black&Blue” clusters was approximatively 50% in BMAT-rich and less than 3% in BMAT-poor sites for sham treated mice (Figure 3e). However, post-VADA treatment, leukemic cells from BMAT-poor sites exhibited a remarkable increase, at least 7 times as compared to Sham treated mice (23% vs. <3% respectively; Figure 3e). These findings show that chemotherapy participates in the selection of similar human T-ALL LSC sub-populations in BMAT-poor and -rich sites.

To further characterize the selected leukemic cells post-VADA treatment, we analyzed CD44 expression of leukemic cells from treated or untreated femurs of mice xenografted with human T-ALL. Leukemic cells from VADA-treated femurs displayed higher CD44 transcript and protein expression compared to Sham-treated condition (Figure 3f-g). As expected from the used cell-cycle targeting drugs (like Vincristine and Cytarabine), leukemic cells from VADA-treated femurs were mainly quiescent (Figure 3h-i).

Together, our data show the presence of leukemic cells in BMAT-poor sites, phenotypically similar to resistant leukemic cells present in BMAT-rich sites, capable of evading chemotherapy. Furthermore, CD44^high^ expression correlates with a quiescent state and is a reliable marker to track chemoresistance.

### *CD44^high^* and *Ki67^neg/low^* biomarkers identify a cell population present in T-ALL patients

To ensure physiological relevance of the prevously described results, we investigated the presence of *CD44^high^ Ki67^neg/low^* leukemic cells from three T-ALL patients by scRNAseq (patient details in Supplementary Table 1), similar to BMAT-rich sites from PDX models. Paired diagnosis/relapse samples were compared (Figure 4 and Supplementary Figure 10). In a 4^th^ library, we analysed 2 additional patients at diagnosis (Supplementary Figure 11). The results of UMAP and clustering analyses of pooled cells from each patient indicated several clusters, but diagnosis and relapse samples had specific separated gene profiles (Figure 4a-b and Supplementary Figure 10a-b; f-g). The same observations were found with the 2 additional T-ALL patients at diagnosis (Supplementary Figure 11a-b). Normal cells, that may contaminate leukemic samples, were excluded by 1) the absence of oncogenic fusion (TLX3 expression in Figure 4c-d) or 2) using the expression of specific lineage genes (B cells : CD79A, CD79B, CD19, MS4A1; NKT cells : NKG7, GNLY and Monocytes : FCGR3A, S100A4, LYZ, CST3, CD33, CD14) (Supplementary Figures 10c, 10h and 11c). Interestingly, expression of *CD44* and *Ki67* were observed in all samples and were mutually exclusive (Figure 4e, Supplementary Figures 10d, 10i and 11d). Focusing on leukemic cells indicated that *Ki67^neg/low^CD44^high^* leukemic cells were only scarcely detected, except for one relapse sample (Figure 4f, Supplementary Figures 10e, 10j and 11e) as in leukemic cells from BMAT-poor sites in T-ALL models (Figure 2h-i). To determine whether *Ki67^neg/low^CD44^high^* leukemic cells found in patients’ samples shared a gene signature with leukemic cells from BMAT-poor sites, we compared their transcriptomic profile (Supplementary Table 6-7-8-9). We selected 38 common significantly upregulated genes obtained from at least 2 libraries (Figure 4g, Supplementary Table 10) of which 24 were found also upregulated in human leukemic cells from Cluster 4 (Figure 1c and Figure 4h, Supplementary Table 11). To determine whether this transcriptomic signature was specific to leukemic cells, we extended our analysis to *Ki67^neg/low^CD44^high^* normal cells found in all libraries (Supplementary Figure 12a-d, Supplementary Table 12-13-14-15). We compared the transcriptomic signature found in *Ki67^neg/low^ CD44^high^* leukemic cells with all significantly upregulated genes found in *Ki67^neg/low^ CD44^high^* normal cells from the 4 libraries (Supplementary Figure 12e, Supplementary Table 16). We found that 10 genes were common, meaning that 14 genes that composed the transcriptomic signature derived from leukemic cells are specifically leukemia-related (Supplementary Figure 12f, Supplementary Table 17). These observations show the presence of rare quiescent CD44^high^ leukemic cells in patient samples that bear a similar expression signature than human T-ALL from chemoresistant and BMAT-poor/rich sites from PDX models. To explore the relevance of high *CD44* expression as a prognosis biomarker, we evaluated *CD44* expression level in a Children’s Oncology Group-Therapeutically Applicable Research to Generate Effective Treatments (COG-TARGET) cohort, in which expression profiles of 264 diagnostic human T-ALL samples are presented^37^. Strikingly, data revealed that patients with high *CD44* mRNA expression were significantly associated with HOXA, LMO2/LYL1 oncogenic subgroups, known to be enriched in early T-cell precursors ALL (ETP-ALL) (Supplementary Figure 13a-b)^37–39^ and to present a poor prognosis after chemotherapy^40^. Together, these results show that quiescent *CD44^high^* cells described in human T-ALL PDX models are also detected in T-ALL patients samples, they have a specific gene signature and *CD44* expression levels are enriched in subgroups that are associated with a poor prognosis.

**Figure 4:**
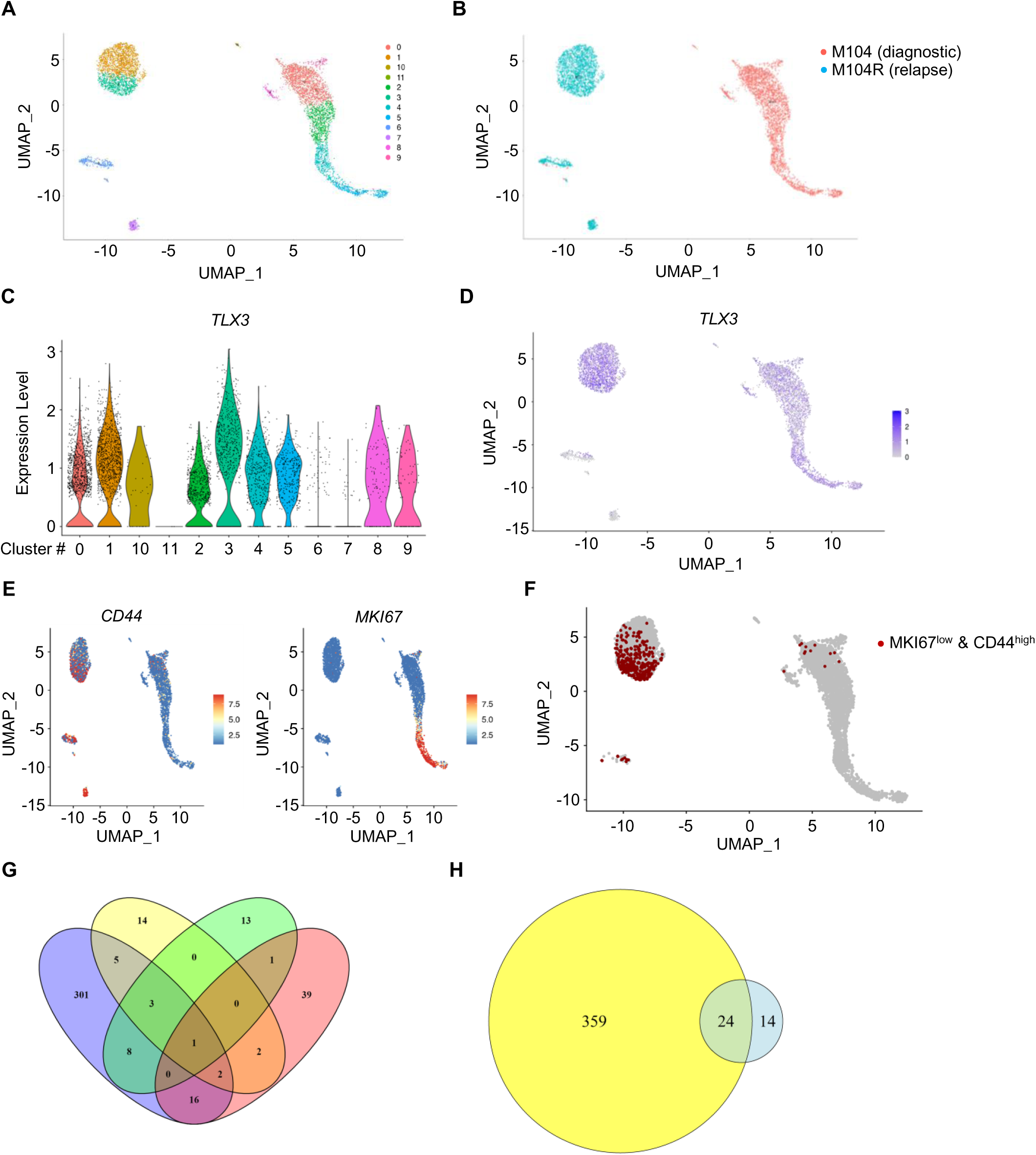
*Ki67^neg/low^ CD44^high^* population is found in diagnosis and relapse T-ALL samples. **A-F** Single Cell RNA sequencing of paired Diagnosis (M104) and Relapse (M104R) Human T-ALL. UMAP color-coded clustering (**A**) and depending of Diagnosis or Relapse origin (**B**). **C-D** *TLX3* Expression level (**c**) overlaid on UMAP representation (**d**). Expression levels of *MKI67* and *CD44* overlaid on UMAP representation (**E**). **F** *MKI67^neg/low^ CD44^high^* population (red dot) overlaid on UMAP restricted to human T-ALL cells. **G** Venn Diagram of significantly upregulated genes identified in *MKI67^neg/low^ CD44^high^* hT-ALL cell population from 4 libraries. Values indicate number of genes. **H** Venn Diagram of common upregulated genes identified in *MKI67^neg/low^ CD44^high^* in at least 2 libraries (38 genes) and significantly upregulates genes identified in Cluster 4 (Fig. 1). Human T-ALL samples are described in Supplementary Table 1.

### CD44^high^ cells display an increased E-selectin binding ability

CD44 is known to interact with a wide variety of ligands, such as hyaluronic acid and E-selectin^41^. At steady state, endothelial cells whithin the BM naturally express the E-selectin^42^. The expression of E-selectin is further increased by inflammatory agents such as TNF-α or IL-1β-mediated activation^43^, both of which are secreted by adipocytes in the BM^44^. To demonstrate E-selectin binding activity, CD44 but also several other ligands, such as P-selectin glycoprotein ligand-1 (PSGL-1/CD162) and Leukosialin (CD43), must be decorated with multiple sialyl Lewis X (sLe^x^) motifs. Upon analysis of the sLe^x^ motif on leukemic cell surface using HECA-452^45^, we observed that leukemic cells expressing high CD44 from BMAT-rich and poor sites are decorated by sLe^x^ motifs (Figure 5a). Subsequently, we evaluated the attachment efficacy of a chimeric soluble human E-selectin-IgG1 fusion protein to leukemic cells recovered from different BM territories. As anticipated, higher leukemic cells from the BMAT-rich site could bind E-selectin compared to BMAT-poor sites (respective median: 7.8 % and 0.4% E-selectin^+^ cells) (Figure 5b). Interestingly, T-ALL cells from the VADA-treated BMAT-poor site also exhibited higher E-selectin-binding levels (median: 4.5% E-selectin^+^ cells) compared to untreated mice (Figure 5b). Using flow cytometry analysis, we observed that whatever the analyzed BM site, binding to E-selectin was detected for cells with the highest CD44 expression (Figure 5c-d). These observations perfectly support the earlier findings obtained with HECA-452 staining.

**Figure 5:**
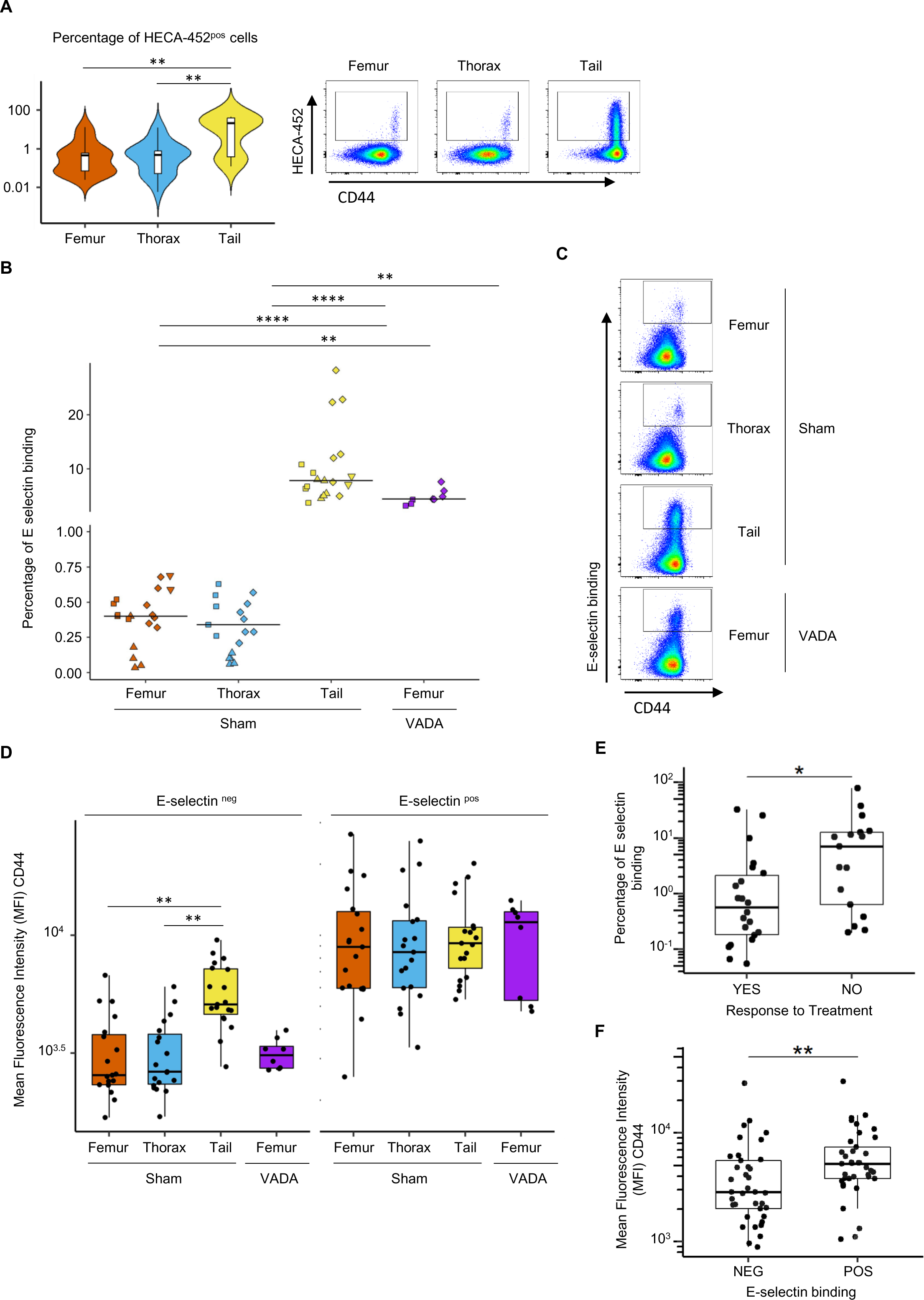
E-selectin binding is enhanced in CD44^high^ population. **A** Percentage of HECA-452 positive cells performed by Flow Cytometry on human CD45^+^/CD7^+^ T-ALL cells from Femur (orange), Thorax (Blue) and Tail Vertebrae (Yellow). PDX of 5 hT-ALL samples (17 mice). Flow Cytometry analysis example obtained with M197-PDX, T-ALL cells unstained (black line), from Femur (orange line), Thorax (blue line) and Tail (yellow line) (inset). **B** Frequency of human CD45^+^/CD7^+^ T-ALL cells from Femur (orange), Thorax (blue), Tail (yellow) of Sham and Femur of VADA mice (purple). **C** Representative E-selectin binding/CD44 staining of human CD45^+^/CD7^+^ T-ALL (M106) cells from Femur, Thorax, Tail of Sham mice and Femur of VADA mice analyzed by Flow Cytometry. **D** MFI of CD44 of human CD45^+^/CD7^+^ T-ALL cells from Femur (orange), Thorax (blue), Tail (yellow) of Sham and Femur of VADA mice (purple) binding or not E-selectin *in vitro*. **E** Proportion of E-selectin binding T-ALL cells according to the Response to Treatment (« NO » vs « YES »). **F** MFI of CD44 expression on human T-ALL cells according to E-selectin binding (« NEG » vs « POS »). **D, E** and **F** Data are shown as box-and-whisker plots of the following numbers of mice for each group. Boxes indicate the 25^th^ and 75^th^ percentiles; whiskers display the range and horizontal lines in each bow represent the median. **A**, **B** and **D** Statistical significance was assessed by Kruskal-Wallis test followed by Dunn’s multiple comparisons test (** p < 0.01; **** p < 0.001). **E** and **F** Statistical significance was assessed by a Mann & Withney test (** p < 0.01). Human T-ALL samples are described in Supplementary Table 1.

E-selectin binding was also measured in leukemic cells from 46 patient samples (Supplementary Figure 14a). Using the clinical information available for 39/46 patients, we could classify the samples into two groups: sensitive and refractory cases, the latter including refractory and relapsed patients. Combining informations of patient treatment response and E-selectin binding levels showed Relapse/Refractory patients (group “NO” response) had ten-fold higher levels of E-selectin binding compared to chemotherapy sensitive patients (group “YES” response) (median 7.0% vs 0.6% (Figure 5e)). Interestingly, regardless of the patient samples analyzed, T-ALL cells more prone to E-selectin binding were CD44^high^ (Figure 5f, Supplementary Figure 14b). Of note, E-selectin binding potential was not related to their EGIL Grade classification for the 29/39 patients for which the information was available (Supplementary Figure 14c). Altogether, we found that T-ALL cells capable of E-selectin binding are CD44^high^ both in T-ALL models and in human patients. From a clinical point of view, our data suggest that E-selectin binding potential stratifies Relapse/Refractory patients.

## Discussion

Although childhood ALL in high-income countries have very high cure rates of around 90%, the prognosis among ALL adults remain poor, with <45% of affected patients expected to achieve long-term disease-free survival. Plus, patients with R/R (Relapse/Refractory) T-ALL face a poor prognosis with Overall Survival (OS) of <10% in adults and <25% in children^3^. In this context, few innovative treatments are proposed. Nelarabine is the only FDA-approved therapy thanks to its demonstrated efficacy in the adult and pediatric populations. Chimeric Antigen Receptor (CAR)-T cell therapy, which reached a success story in B-ALL, started to be translated to T-ALL in the second line after therapeutic failure^46–48^. Obviously, additional research work is necessary to propose novel therapeutic strategies. Interestingly, we and others have observed that ALL cells are able to downregulate their surface marker expression in relation with a decreased translational activity that might contribute to potential CAR-T cell escape already observed in B-ALL^14,22^. In the present work, we demonstrate that the exploration of leukemic cell diversity within BM revealed multiple clusters, with distinct subpopulations residing in adipocyte-rich (BMAT-rich) and –poor (BMAT-poor) sites. We identified a Quiescent cluster, Cluster 4, predominantly composed by leukemic cells from BMAT-rich site, displaying additional cell adhesion properties which implicate CD44. The correlation between high CD44 expression and a Quiescent state suggests a potential marker for tracking chemoresistant cells which is supported by the fact that CD44 is also associated with chemoresistance in different hematological diseases^25,26,28^. We show that transcriptional and translational upregulation of CD44 is associated to sLe^x^ motif adding and, as expected, an advantage to E-selectin binding. Of note, fucose addition mediated by a variety of FucosylTransferase (FUT3/4/5/6/7) in α(1,3)-linkage to terminal sialylactosamines on CD44/CD162/CD43 creates E-selectin ligands^24,49,50^. The single cell transcriptomic analysis shows that only quiescent *CD44*^high^ T-ALL cells possess *FUT7* expression (data not shown), which participates to the sLe^x^ motif creation and allows the E-selectin binding for the Ki67^neg/low^CD44^high^ LSC population. On note, other studies have shown that FUTs expression may be a prognostic marker associated with bad clinical outcomes in different tumors^51–55^. Here in T-ALL cells, we observe that LSC cells, more or less numerous in respectively BMAT-rich and -poor BM, are able to bind E-selectin in experimental models, very similar to cells being also detected in patients. It is reported that pro-inflammatory cytokines (TNF-α and IL-1β) enhance E-selectine expression in inflammed BM endothelium^42,56^. Associated with the fuel energy role of adipose tissue, adipocytokines, that include TNF-α and IL-1β which are highly expressed in BMAT^44^, must now be taken into consideration for their role in the microenvironment evolution after chemo/radiotherapy and aging^11,13,14,57^. We speculate that E-selectin expressed by endothelial cells, especially in BMAT-rich site, may attach Ki67^neg/low^ CD44^high^ T-ALL cells and thus constitute a chemoresistant BM niche. Uproleselan (developed by the Glycomimetics company), which disrupt E-selectin receptor/ligand interactions is being evaluated in a Phase 3 trial for the treatment of Relapsed/Refractory AML^58,59^. This antagonist could represent a new and original therapeutic molecule to target quiescent chemoresistant T-ALL cells.

Even though recent results reveal that chemoresistant T-ALL cells are not located in a specific niche, they are all consistent with the observations that the quiescent state is a specific behavior associated with leukemic cell chemoresistance^36,60^. Our results obtained with *in vivo* VADA treatment support this conclusion. Moreover, we demonstrate that the quiescent state observed after treatment in BMAT-poor sites resembles the quiescent state observed in BMAT-rich sites, with an intense CD44 expression (CD44^high^) and increased E-selectin binding. These observations support the idea that the Ki67^neg/low^CD44^high^ leukemic cells endowed with E-selectin binding define a chemoresistant leukemic cell population.

Relapse remains the major cause of death in ALL. Ebinger *et al* described a chemoresistance model based on the persistence of dormant/non-dividing BCP-ALL cells (called Label-Retaining Cells, LRCs) located close to BM endosteal regions. LRCs display a transcriptomic signature that is enriched in MRD compared to diagnosis^23^. In our work, the scRNAseq Cluster 4, in which we identified Ki67^neg/low^CD44^high^ cells, is highly enriched for the LRC signature, indicating common chemoresistance phenotypes. Moreover combined scRNAseq of leukemic cells from patients and from experimental models precisely report that Ki67^neg/low^ CD44^high^ leukemic cells harbor a specific 14 genes-transcriptomic signature. In AML, a 17 or 6 genes score has been defined based on gene expression of Leukemic Stem Cells (LSCs) endowed with properties like dormancy, and is providing strong prognostic information^61,62^. Similarly, a 9-genes score is proposed for helping risk stratification in B-ALL, based on the LRCs transcriptomic signature^63^. Among the 14 transcripts of our Ki67^neg/low^CD44^high^ T-ALL cell signature, some encode for markers implicated in cancer cell migration (MALAT1, Metastasis-associated lung adenocarcinoma transcript 1^64^), in Immune Checkpoint / Tolerance (HLA-B/C/E, Human Leukocyte Antigen^65^), in protein synthesis through the inhibition of rRNA processing (PNRC1, Proline-rich Nuclear Receptor Coactivator 1^66^) and drug resistance (ARL4C, ADP ribosylation like factor 4C^67^). Interestingly, MALAT1 was significantly upregulated in MRD+ compared to MRD-patients and highly expressed in relapsed compared to new case patients of pediatric T- and B-ALL^68^. Our 14 genes-signature might be considered as a new MRD signature specific to T-ALL especially since 9 of them are also present in the B-ALL LRC signature^23^.

In summary, this study provides comprehensive insights into the heterogeneity of T-ALL cells within the BM microenvironment and identifies a specific transcriptomic signature of T-ALL chemoresistance. The expression of the sLe^x^ motif, which allows the binding to E-selectin, associated with high CD44 expression highlights its potential as a prognostic biomarker of chemoresistance. Our data further demonstrate that E-selectin binding is a relevant marker for Relapse/Refractory stratification.

## Materials and methods

### Human T-ALL

Blood and/or bone marrow samples from patients with human (hu)T-ALL were collected at Hôpital R. Debré, Hôpital A. Trousseau (Paris, France) and Hôpitaux Civils de Lyon (Lyon, France) and processed as previously described^22^. Newly diagnosed leukemic cells was partly used directly after sampling for *in vitro* and *in vivo* experiments and partly frozen in Foetal Calf Serum (FCS) containing 10% of Dimethylsufoxide (DMSO) (D8418, Sigma-Aldrich). Human T-ALL samples are described in Supplementary Table 1.

### Human T-ALL xenografts

Patient derived xenografts (PDX) were established from T-ALL biopsy samples in nonobese diabetic/severe combined immunodeficiency / interleukin-2Rγ null mice (NSG, The Jackson Laboratory, Bar Harbor, USA) that are produced in pathogen-free animal facilities (Commissariat à l’Energie Atomique et aux Energies Alternatives [CEA], Fontenay-aux-Roses, France). Leukemic cells were injected intravenously in non-irradiated mice. Bone marrow (BM) samplings, performed after anesthesia (Isoflurane) and analgesia (3µg/mL Buprenorphine), allowed to monitor human T-ALL expansion *in vivo*. Leukemic cell infiltration was evaluated using immuno-labeling with anti-huCD45 and anti-huCD7 antibodies (see Supplementary Table 2) and a FACS-Canto II flow cytometer (Becton Dickinson, BD). Cells from Femur, Thoracic or Tail Vertebrae were isolated^22^. For RT-PCR and scRNAseq analysis, human T-ALL cells were purified after labeling with anti-huCD45 PE conjugated antibody followed by immuno magnetic selection with anti-Phycoerythrin (PE) microbeads (130-048-801, Miltenyi Biotec).

### Flow cytometry

Phenotype, cell cycle progression and absolute number of human T-ALL cells, were measured by flow cytometry using FACS Canto II and LSR II apparatus (BD). Data analysis was carried out with FlowJo software. Cells were stained with fluorescein (FITC)-, phycoerythrin (PE)-, Peridinin chlorophyl protein-Cyanine 5.5 (PerCP Cy5.5)-, PE-cyanin7 (PC7)-, allophycocyanin (APC)-, allophycocyanin eFluor780 (APC-eFluor780)-, allophycocyanin Vio770 (APC-Vio770)-, Brillant Violet 421 (BV421)-conjugated mouse monoclonal antibodies specific for human markers. All antibodies were purchased from BD Pharmingen, Miltenyi Biotec or e-Bioscience (Supplementary Table 2). Absolute number of cells was quantified by determining the number of huCD45^+^/huCD7^+^ cells in BM sampling. For cell cycle analysis, cells were staining with anti-huCD45 and anti-huCD7 antibodies then permeabilized using Cytofix/Cytoperm (554722, BD Biosciences) during 15 minutes at 4°C, washed with Perm/Wash Buffer (554723, BD Biosciences) then labeled with anti-Ki67 antibodies (556027 or 556026, BD Biosciences) at 4°C during 45 minutes. Hoechst 33342 (H3570, Life Technologies) is added at 20µg/mL final 10 minutes before the end of incubation. Cells were washed with Perm/Wash Buffer and resuspended in PBS.

### 10x genomics single-cell package and libraries preparation

After purification, cells are counted with trypan blue then suspended at 1 x 10^6^ cells/mL in PBS 0.04% BSA. 20 000 cells per condition were loaded in Chromium Next GEM Chip G. Single-cell libraries were generated using Chromium Next GEM Single Cell 3’ Reagent Kits (v3.1): GEM, Library and Gel Bead Kit v3.1 (PN-1000128), Chip G Single Cell kit (PN-1000127) and Dual Index kit TT Set A (PN-1000215) (10x Genomics), and following the Chromium Next GEM Single Cell 3’ Reagent Kits (v3.1) User Guide (manual part no. CG0000204 Rev D). Reverse transcription and libraries preparation were performed on Veriti 96-Well Thermal Cycler (ThermoFisher Scientific). Amplified cDNA and libraries were verified on Bioanalyzer 2100 (Agilent Technologies) using High Sensitivity DNA kit (Agilent Technologies). Libraries were sequenced (Read1: 28bp, Index7: 10bp, Index5: 10bp and Read2: 90bp) on NovaSeq 6000 Sequencing System (Illumina).

### Pre-processing of scRN-seq data

Sequencing results were demultiplexed then converted to FASTQ format. Galaxy was used to process raw data, demultiplexing, barcode processing, and alignment on GRCh38/hg38 reference genome. EmptyDroplet method was used to remove Barcodes that are associated to very low amount of UMIs (< 50 UMIs). After this process, we obtained 11334, 10914 and 11356 cells in respectively Femur, Thoracic and Tail Vertebrae. Quality Filtering, Normalization, Dimensionality Reduction, Unsupervised Clustering and Find the differentially expressed genes analysis were performed using the *Seurat v4* R package^69^. Filtration/pre-processing was done to remove low quality of cells. Indeed, cells with more than 10% of mitochondrial transcripts were removed and in which there are less than 100, 95 or 85 mRNA respectively in Femur, Thoracic and Tail vertebrae (Supplementary Figure 1B). After pre-processing, the median of detected genes and transcripts per cell were 2113 and 4978 respectively (Figure 1A). Data was normalized by global median mRNA account and log-transformed.

### Dimensionality Reduction

For each dataset, we identified a subset of 2000 variable genes with the highest dispersion using *FindVariableFeatures* function of *Seurat v4* package^69^. Dimensionality reduction was done using these variable genes. In order to remove batch effect, we use the RCPA (Reciprocal PCA) method. First we find a set of anchors between the three Seurat Objects (Femur, Thoracic and Tail Vertebrae) thanks to FindIntegrationAnchors function and then, we integrate the objects using *IntegrateData* function. It generates a batch corrected expression matrix used as input for Principal Component Analysis (PCA). We identified tha first 35 principal components as relevant based on the Jack Straw method (JackStraw and ScoreJackStraw functions) by looking at a p-value drop off of principal components observed on *Jack Straw Plot*.

### Visualization and Clustering

To visualize the data, we reduced the dimensionality to project cells in 2D space using *RunUMAP (Uniform Manifold Approximation and Projection)* function^70^. Clusters were identified with *FindClusters* function following *FindNeighbors* function with resolution, res = 0.4. Whatever the resolution used, Cluster 4 keep his identity up to res = 1.2 (Data not shown).

### Differential expression of genes signatures

In order to find differentially expressed genes per cluster we performed the Wilcoxon Rank Sum test from *FindAllMarkers* function. The significance threshold was p-value adjusted (Bonferroni correction) < 0.05 and log2FoldChange > 0.25.

### *In vivo* chemotherapy

Conventional drug treatment was adapted from Samuels et al, 2014. When the %huT-ALL in BM reached at least 70%, mice were randomized for chemotherapy treatment. We injected a 1-week schedule of Vincristine (0.25mg/kg, i.v., Monday), cytarabine (Ara-C, 2.5mg/kg, i.v., Monday), Dexamethasone (5mg/kg, i.p., each day Monday-Friday) and L-Asparaginase (1000U/kg, i.p., each day Monday-Friday), further called VADA. During treatment, mice were closely monitored for signs of drug-related toxicity (weight-loss [Supplementary Figure 7B], lethargy, shaggy hair) and euthanized at the first sign of morbidity. At the end of treatment (3 days after the last injection), mice are euthanized and the BM infiltration was evaluated.

### RNA extraction and Real-time PCR

Total RNA was extracted from T-ALL cells collected purification using RNeasy Plus Mini or Micro Kit (Qiagen). cDNA were generated using SuperScript® VILO^TM^ cDNA Synthesis Kit (ThermoFisher Scientific).Real-time PCR reactions were carried out using *Power* SYBR® Green PCR Master Mix (ThermoFisher Scientific) and run on StepOne Real-time PCR system (Applied Biosystems). Forward and reverse primer sequences are described in Supplementary Table 3. Datas were normalized over *GAPDH* Ct values.

### Western Blot

Proteins were extracted with lysis buffer containing 100mM Tris pH8, 100mM NaCl, 1mMEDTA, 1mM EGTA, 1% NP40, 0,5% DOC, 50% Glycerol, 0,1% SDS and cocktail of protease inhibitors, separated by 4-12% SDS PAGE, transfered onto nitrocellulose membrane (Schleicher & Schuell) and immunoblotted in standard conditions. The primary antibodies used were: mouse anti-human CD44s pan specific and rabbit anti-b-actin; secondary antibodies were: goat anti-mouse and goat anti-rabbit. All antibodies were purchased from Bio-Techne, Sigma-Aldrich or Abcam (Supplementary Table 2)

### E-Selectin Binding

For analysis of E-selectin binding potential, recombinant E-selectin– human-IgG1 fusion protein (724 ES 100, R&D Systems) was pre-complexed at 5µg/mL with PE-conjugated goat anti-Human IgG diluted at 1/100^e^ (Supplementary Table 2) for 2 h at room temperature in PBS CaCl_2_ MgCl_2_ (14040133, ThermoFisher Scientific). 200 000 T-ALL cells pre-stained with cell surface markers were incubated with this complexe. After 30 min at room temperature, cells were washed in PBS and analyzed by flow cytometry. For each analysis, a negative (non-binding) control was included, an identical parallel stain in presence of 15 mM EDTA (E-selectin binding is strictly Ca^2+^-dependent).

### Statistical analyses

Statistical analysis was performed using R software. Values were presented as the mean ± SEM (Standard Error of Mean) or median through boxplot. Differences between the means of two experimental conditions and in absence of normal distribution were analyzed using non-parametric Wilcoxon-Mann-Whitney test. For multiple-group comparisons, when n < 30 and in absence of normal distribution, a Kruskal-Wallis test with Dunn’s post test for pairwise multiple comparisons was applied. False Discovery Rates (FDR) were corrected by the Benjamini-Hochberg stepwise adjustment^71^. Differences with p < 0.05 (*), p < 0.01 (**), p < 0.001 (***) or p < 0.0001 (****) are considered statistically significant.

### Study approval

All animal experimentations were conducted after approval of a local ethical committee, and authorization from the French Ministère de l’Enseignement Supérieur et de la Recherche. Informed consent of patients or of relatives was obtained in accordance with the Declaration of Helsinki and the Ethic regulations. The research project has been approved by the ethics evaluation committee of Inserm (IORG0003254, FWA00005831). No compensation was provided to patients.

## Author Contributions

JC and FP conceptualized and supervised the study. JC, BG, IN and BU perfomed mouse experiments. JC, AK and JFP were responsible for analysis of single cell RNA sequencing. JC and FP analyzed data. AP, HL, PB provided human T-ALL samples. JC, TM, SJCM, JFP and FP wrote the original draft of the manuscript. FP provided fundings.

## Acknowledgements

Mouse work was facilitated by Silvia Vincent-Naulleau, Caroline Devanand and Vilma Barroca from the Institut de Radiobologie Cellulaire et Moléculaire, CEA, Fontenay-aux-Roses. Flow Cytometry work was facilitated by Nathalie Dechamps and Jan Baijer. The authors thank Emmanuelle Clappier and Rathana Kim from Hematology Laboratory, Saint Louis Hospital, Paris, France for providing B-ALL samples. The authors also thank Thomas Belmas, Raphael Nicolas for their technical help in experiments.

This work was supported by INSERM, CEA, Université Paris Cité, Université Paris Sud, the Association Laurette Fugain, Fondation LNCC (Equipe labellisée) and the Association The Hope of Princesse Manon.

