## Supplemental Figures for "High CD44 expression identifies rare chemoresistant leukemic cells endowed with enhanced E-Selectin binding in T-ALL"

### Slide 1
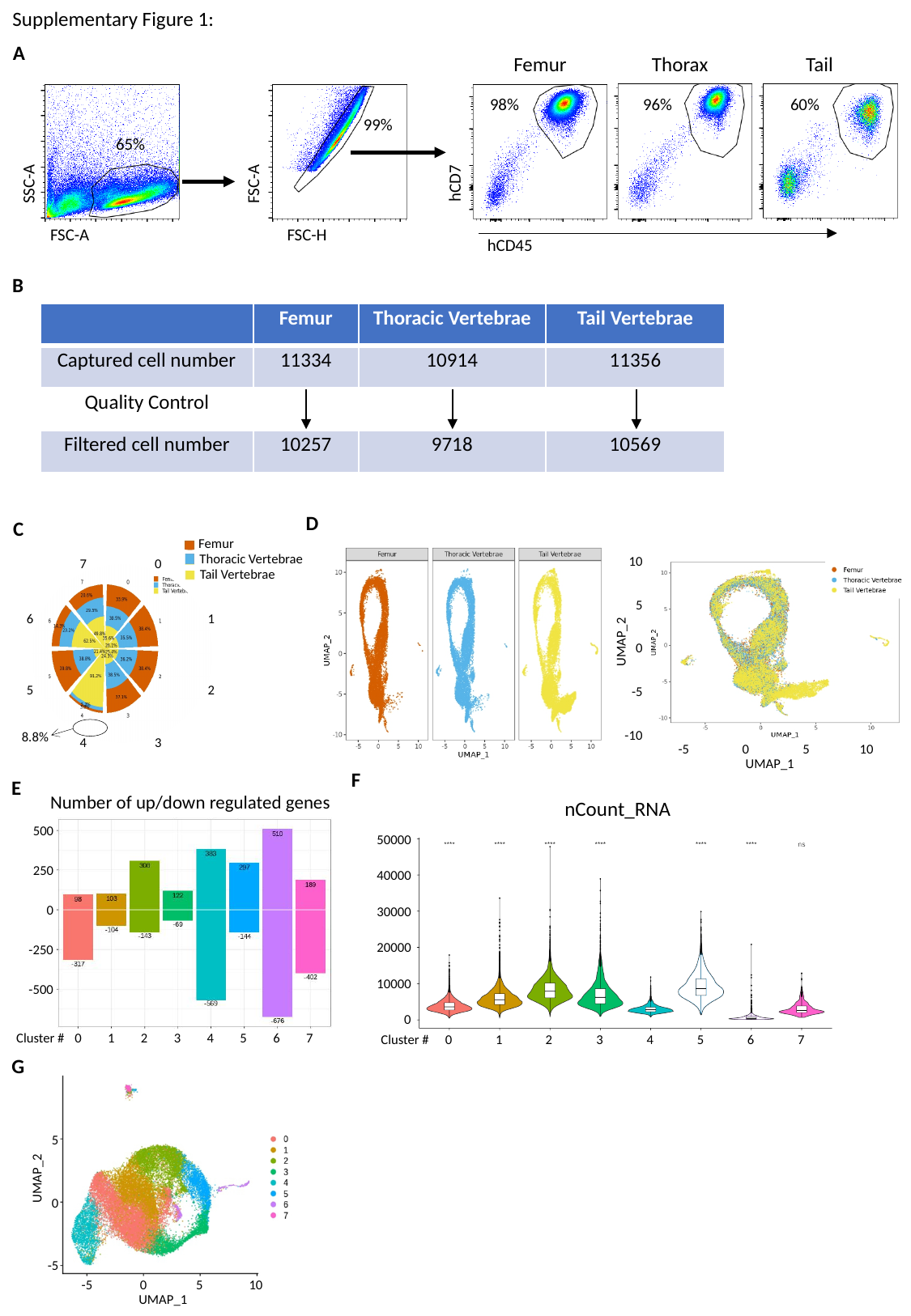

Supplementary Figure 1:
A
Femur
Thorax
Tail
98%
96%
60%
99%
65%
hCD7
FSC-A
SSC-A
FSC-A
FSC-H
hCD45
B
| | Femur | Thoracic Vertebrae | Tail Vertebrae |
| --- | --- | --- | --- |
| Captured cell number | 11334 | 10914 | 11356 |
| Quality Control | | | |
| Filtered cell number | 10257 | 9718 | 10569 |
D
C
7
0
6
1
5
2
8.8%
4
3
Femur
Thoracic Vertebrae
Tail Vertebrae
10
5
UMAP_2
0
-5
-10
-5
0
5
10
UMAP_1
F
E
Cluster #
0
1
2
3
4
5
6
7
Number of up/down regulated genes
500
250
0
-250
-500
nCount_RNA
50000
40000
30000
20000
10000
0
0
1
2
3
4
5
6
7
Cluster #
G
5
UMAP_2
0
-5
0
5
10
-5
UMAP_1

### Slide 2
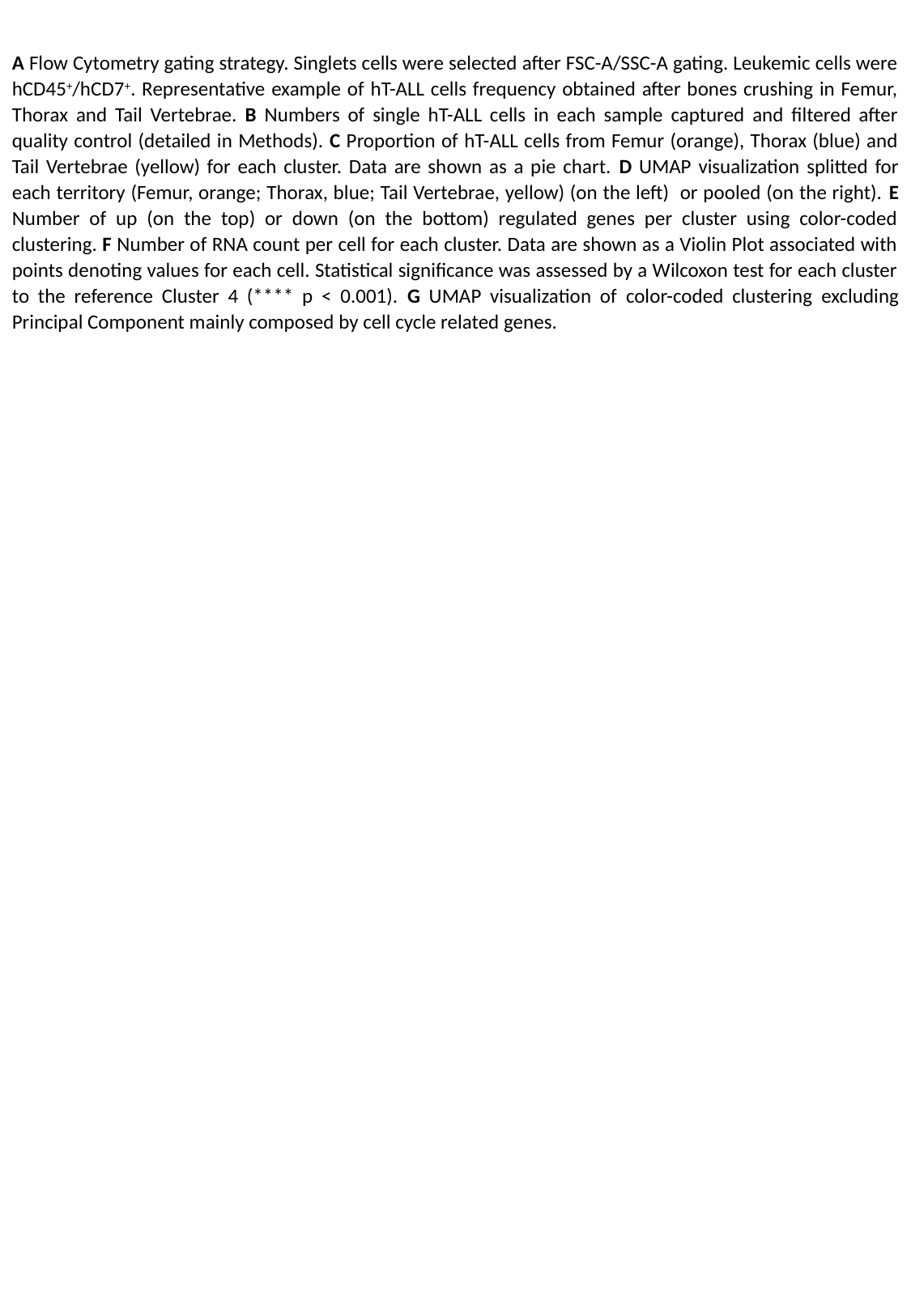

A Flow Cytometry gating strategy. Singlets cells were selected after FSC-A/SSC-A gating. Leukemic cells were hCD45^+^/hCD7^+^. Representative example of hT-ALL cells frequency obtained after bones crushing in Femur, Thorax and Tail Vertebrae. B Numbers of single hT-ALL cells in each sample captured and filtered after quality control (detailed in Methods). C Proportion of hT-ALL cells from Femur (orange), Thorax (blue) and Tail Vertebrae (yellow) for each cluster. Data are shown as a pie chart. D UMAP visualization splitted for each territory (Femur, orange; Thorax, blue; Tail Vertebrae, yellow) (on the left) or pooled (on the right). E Number of up (on the top) or down (on the bottom) regulated genes per cluster using color-coded clustering. F Number of RNA count per cell for each cluster. Data are shown as a Violin Plot associated with points denoting values for each cell. Statistical significance was assessed by a Wilcoxon test for each cluster to the reference Cluster 4 (**** p < 0.001). G UMAP visualization of color-coded clustering excluding Principal Component mainly composed by cell cycle related genes.

### Slide 3
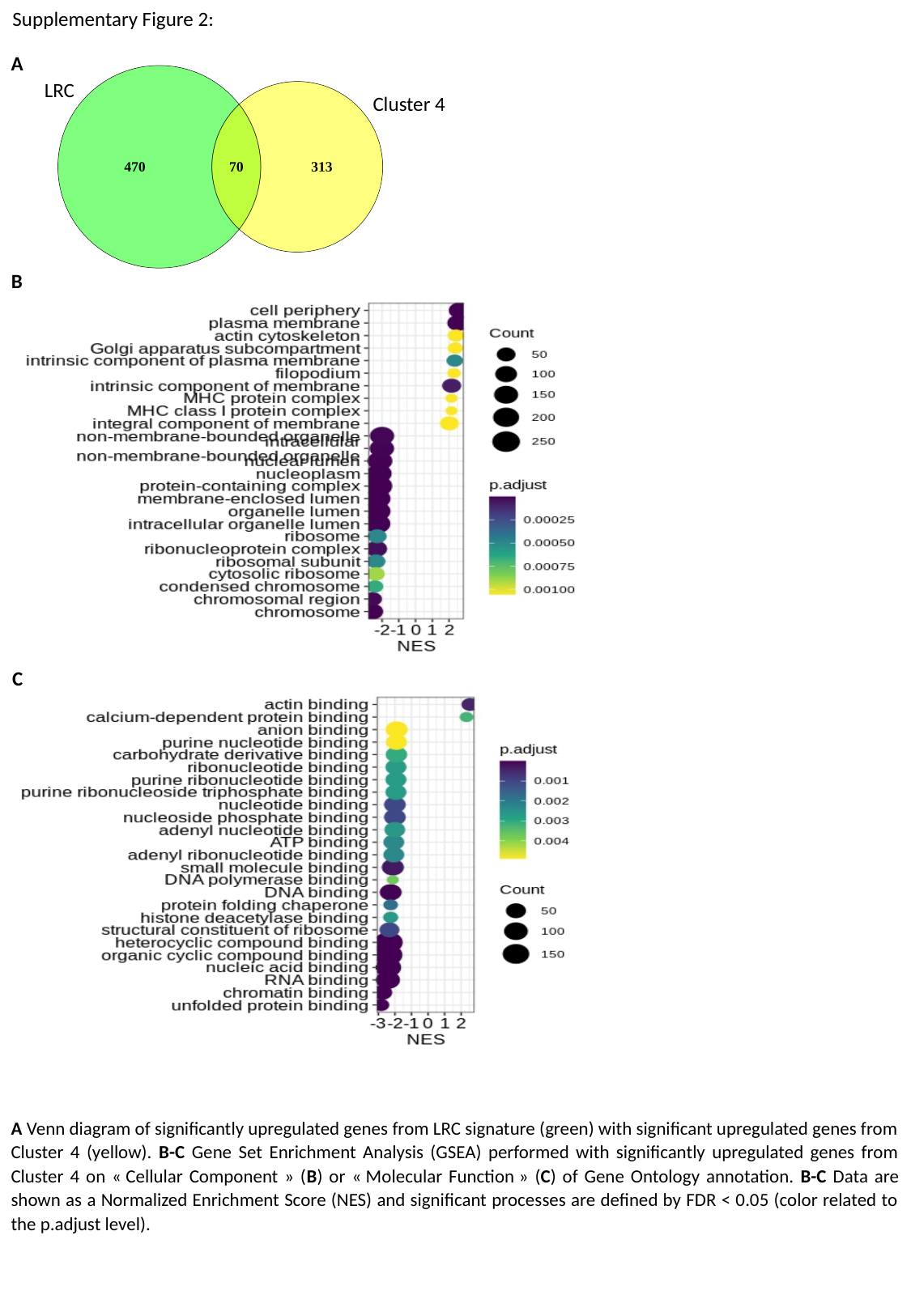

Supplementary Figure 2:
A
LRC
Cluster 4
B
C
A Venn diagram of significantly upregulated genes from LRC signature (green) with significant upregulated genes from Cluster 4 (yellow). B-C Gene Set Enrichment Analysis (GSEA) performed with significantly upregulated genes from Cluster 4 on « Cellular Component » (B) or « Molecular Function » (C) of Gene Ontology annotation. B-C Data are shown as a Normalized Enrichment Score (NES) and significant processes are defined by FDR < 0.05 (color related to the p.adjust level).

### Slide 4
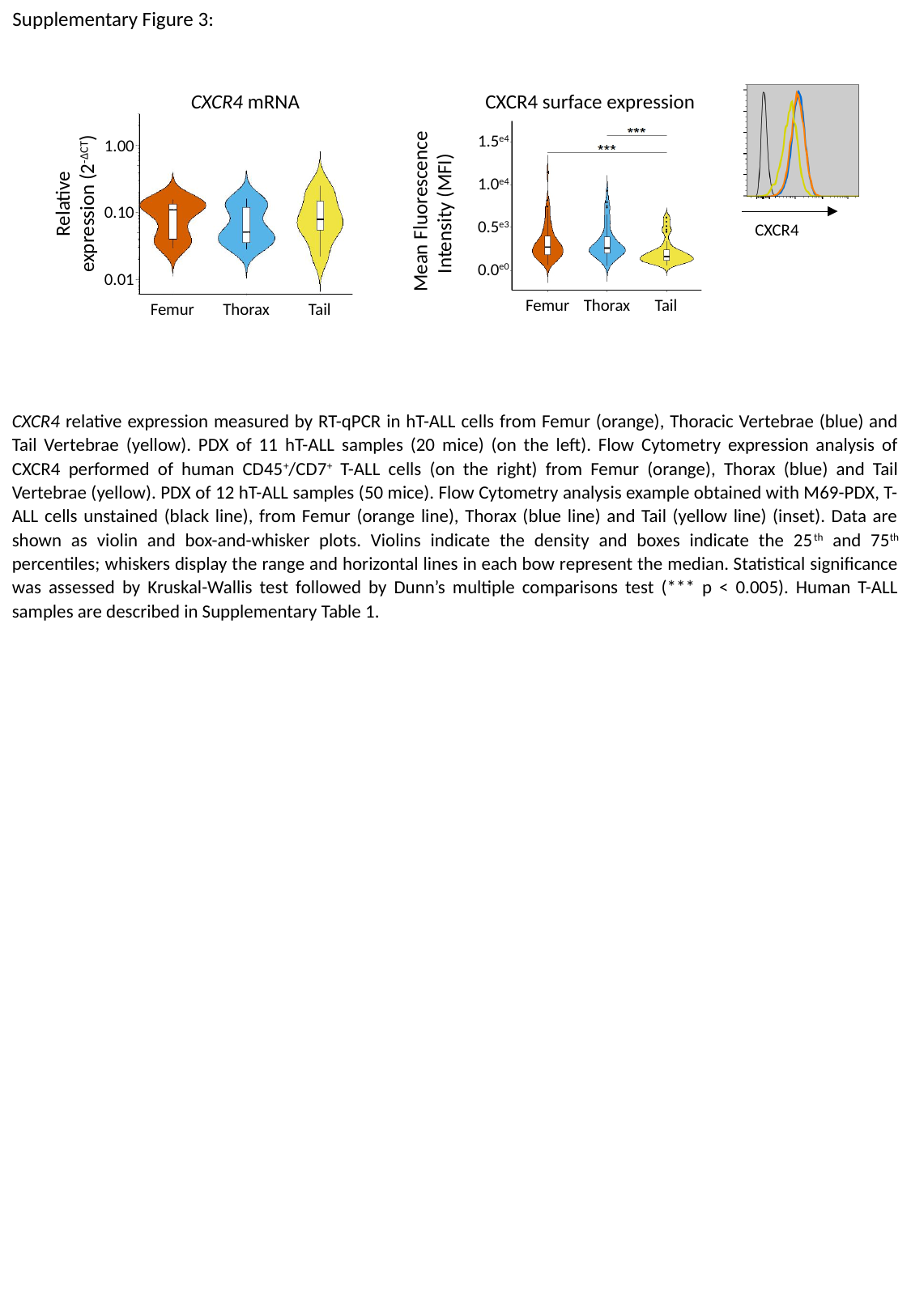

Supplementary Figure 3:
CXCR4 surface expression
Mean Fluorescence Intensity (MFI)
CXCR4
1.5^e4^
1.0^e4^
0.5^e3^
0.0^e0^
Femur
Thorax
Tail
CXCR4 mRNA
Relative expression (2^-ΔCT^)
1.00
0.10
0.01
Femur
Thorax
Tail
CXCR4 relative expression measured by RT-qPCR in hT-ALL cells from Femur (orange), Thoracic Vertebrae (blue) and Tail Vertebrae (yellow). PDX of 11 hT-ALL samples (20 mice) (on the left). Flow Cytometry expression analysis of CXCR4 performed of human CD45^+^/CD7^+^ T-ALL cells (on the right) from Femur (orange), Thorax (blue) and Tail Vertebrae (yellow). PDX of 12 hT-ALL samples (50 mice). Flow Cytometry analysis example obtained with M69-PDX, T-ALL cells unstained (black line), from Femur (orange line), Thorax (blue line) and Tail (yellow line) (inset). Data are shown as violin and box-and-whisker plots. Violins indicate the density and boxes indicate the 25^th^ and 75^th^ percentiles; whiskers display the range and horizontal lines in each bow represent the median. Statistical significance was assessed by Kruskal-Wallis test followed by Dunn’s multiple comparisons test (*** p < 0.005). Human T-ALL samples are described in Supplementary Table 1.

### Slide 5
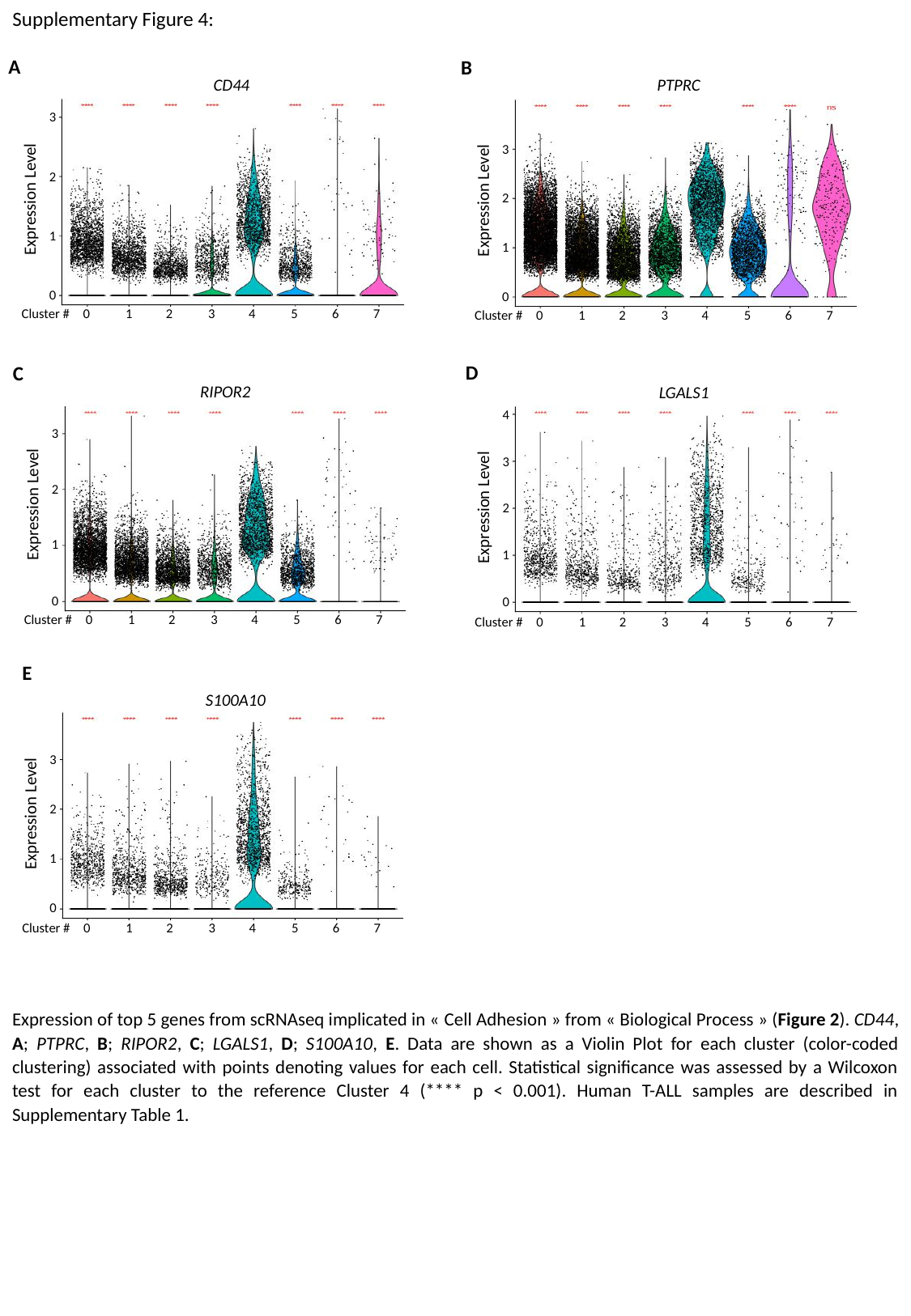

Supplementary Figure 4:
A
CD44
3
2
Expression Level
1
0
Cluster #
1
4
6
7
0
2
3
5
B
PTPRC
3
2
Expression Level
1
0
Cluster #
1
4
6
7
0
2
3
5
D
LGALS1
4
3
Expression Level
2
1
0
Cluster #
1
4
6
7
0
2
3
5
C
RIPOR2
3
2
Expression Level
1
0
Cluster #
1
4
6
7
0
2
3
5
E
S100A10
3
2
Expression Level
1
0
Cluster #
1
4
6
7
0
2
3
5
Expression of top 5 genes from scRNAseq implicated in « Cell Adhesion » from « Biological Process » (Figure 2). CD44, A; PTPRC, B; RIPOR2, C; LGALS1, D; S100A10, E. Data are shown as a Violin Plot for each cluster (color-coded clustering) associated with points denoting values for each cell. Statistical significance was assessed by a Wilcoxon test for each cluster to the reference Cluster 4 (**** p < 0.001). Human T-ALL samples are described in Supplementary Table 1.

### Slide 6
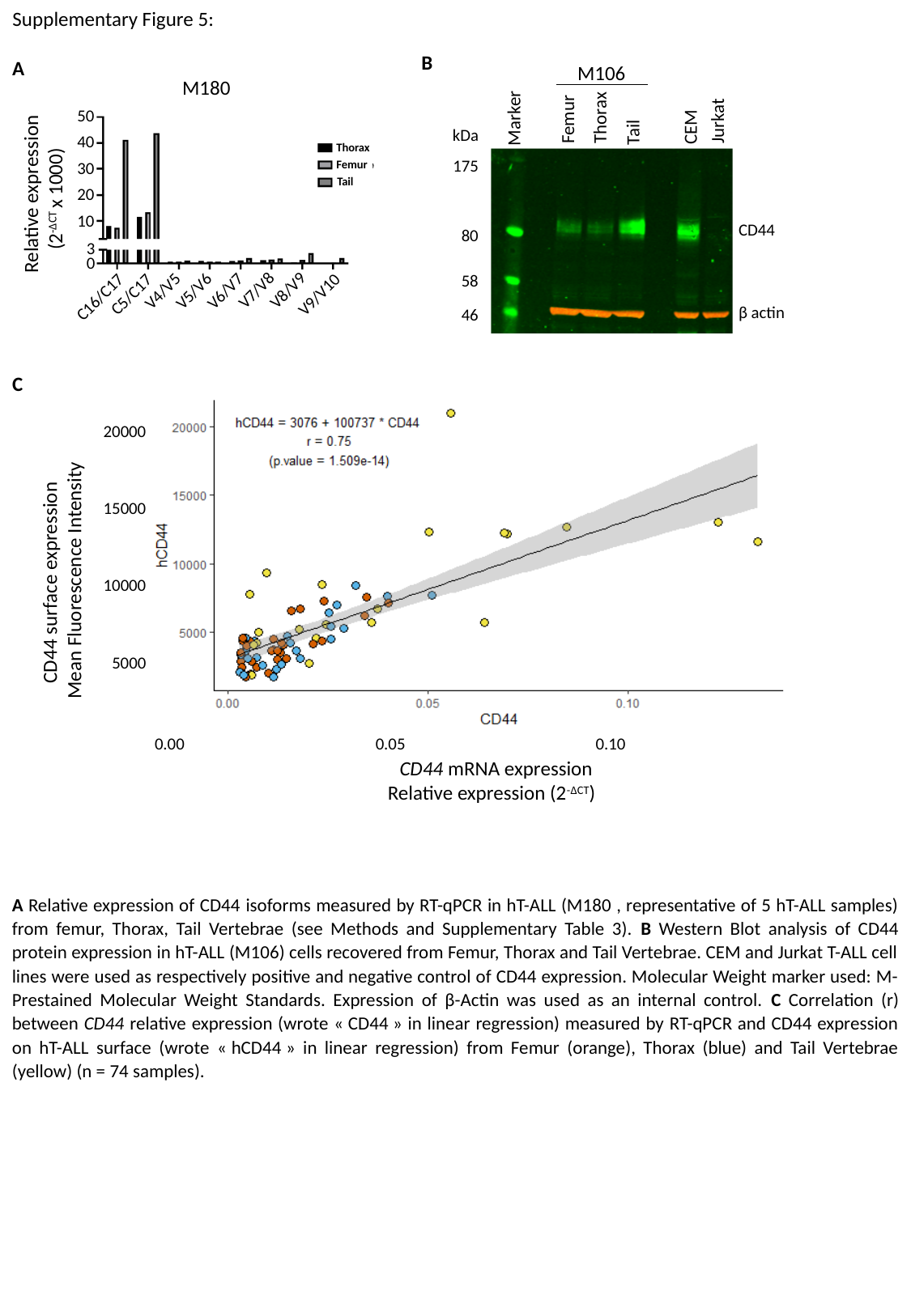

Supplementary Figure 5:
B
Thorax
Marker
Femur
Jurkat
CEM
Tail
kDa
175
80
58
46
CD44
β actin
M106
A
M180
50
40
30
Relative expression (2^-ΔCT^ x 1000)
20
10
3
0
V7/V8
V8/V9
V4/V5
V5/V6
V6/V7
C5/C17
V9/V10
C16/C17
Thorax
Femur
Tail
C
CD44 surface expression
Mean Fluorescence Intensity
CD44 mRNA expression
Relative expression (2^-ΔCT^)
0.00
0.05
0.10
20000
15000
10000
5000
A Relative expression of CD44 isoforms measured by RT-qPCR in hT-ALL (M180 , representative of 5 hT-ALL samples) from femur, Thorax, Tail Vertebrae (see Methods and Supplementary Table 3). B Western Blot analysis of CD44 protein expression in hT-ALL (M106) cells recovered from Femur, Thorax and Tail Vertebrae. CEM and Jurkat T-ALL cell lines were used as respectively positive and negative control of CD44 expression. Molecular Weight marker used: M-Prestained Molecular Weight Standards. Expression of β-Actin was used as an internal control. C Correlation (r) between CD44 relative expression (wrote « CD44 » in linear regression) measured by RT-qPCR and CD44 expression on hT-ALL surface (wrote « hCD44 » in linear regression) from Femur (orange), Thorax (blue) and Tail Vertebrae (yellow) (n = 74 samples).

### Slide 7
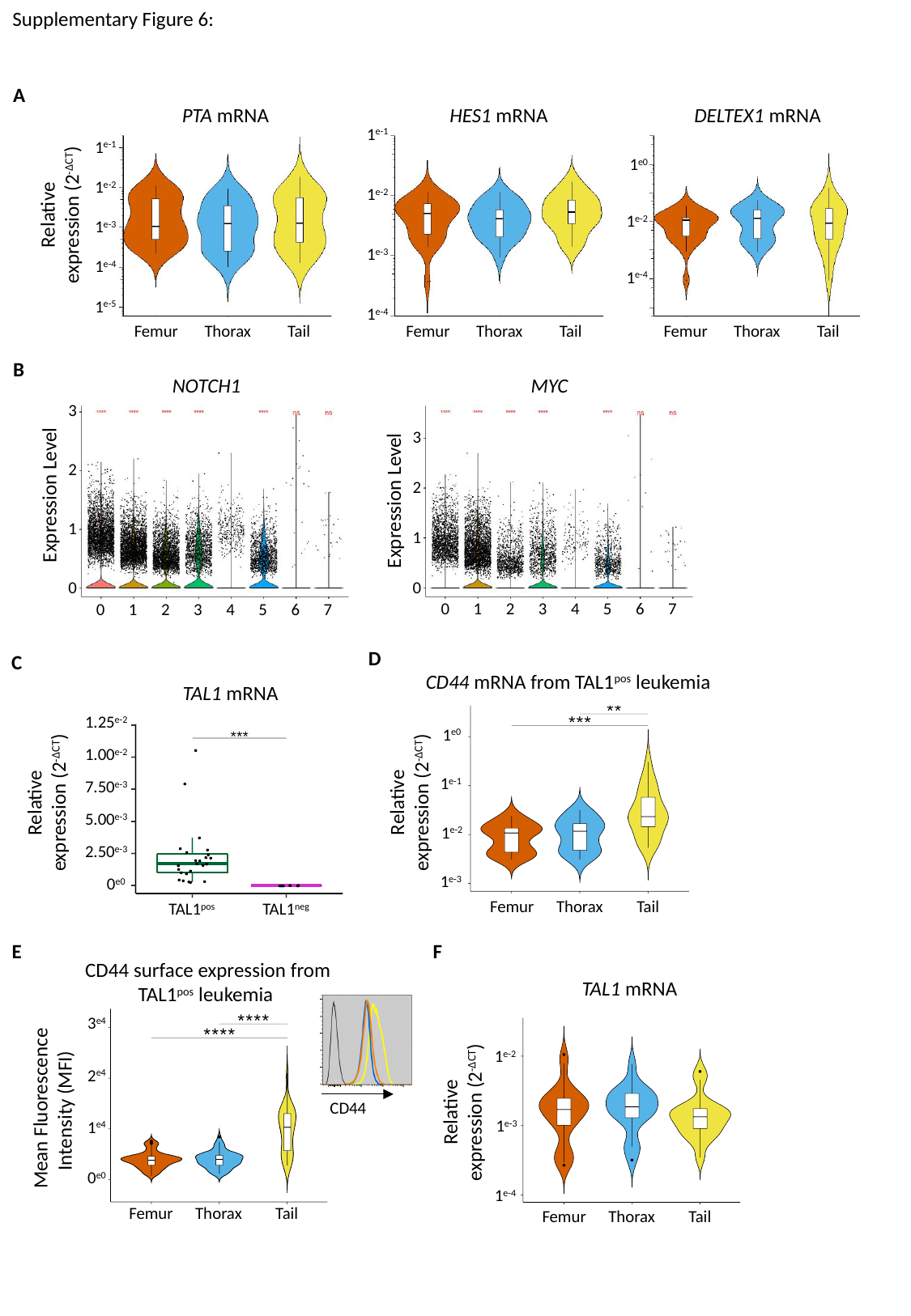

Supplementary Figure 6:
A
PTA mRNA
HES1 mRNA
DELTEX1 mRNA
Relative expression (2^-ΔCT^)
B
Femur
Thorax
Tail
Femur
Thorax
Tail
Femur
Thorax
Tail
1^e-1^
1^e-2^
1^e-3^
1^e-4^
1^e-5^
1^e-1^
1^e-2^
1^e-3^
1^e-4^
1^e0^
1^e-2^
1^e-4^
NOTCH1
MYC
0
1
2
3
4
5
6
7
0
1
2
3
4
5
6
7
3
2
Expression Level
1
0
3
2
Expression Level
1
0
D
Femur
Thorax
Tail
1^e0^
1^e-1^
1^e-2^
1^e-3^
Relative expression (2^-ΔCT^)
C
TAL1 mRNA
Relative expression (2^-ΔCT^)
1.25^e-2^
1.00^e-2^
7.50^e-3^
5.00^e-3^
2.50^e-3^
0^e0^
TAL1^pos^
TAL1^neg^
CD44 mRNA from TAL1^pos^ leukemia
F
1^e-2^
1^e-3^
1^e-4^
Femur
Thorax
Tail
Relative expression (2^-ΔCT^)
TAL1 mRNA
E
CD44
3^e4^
2^e4^
1^e4^
0^e0^
Femur
Thorax
Tail
Mean Fluorescence Intensity (MFI)
CD44 surface expression from TAL1^pos^ leukemia

### Slide 8
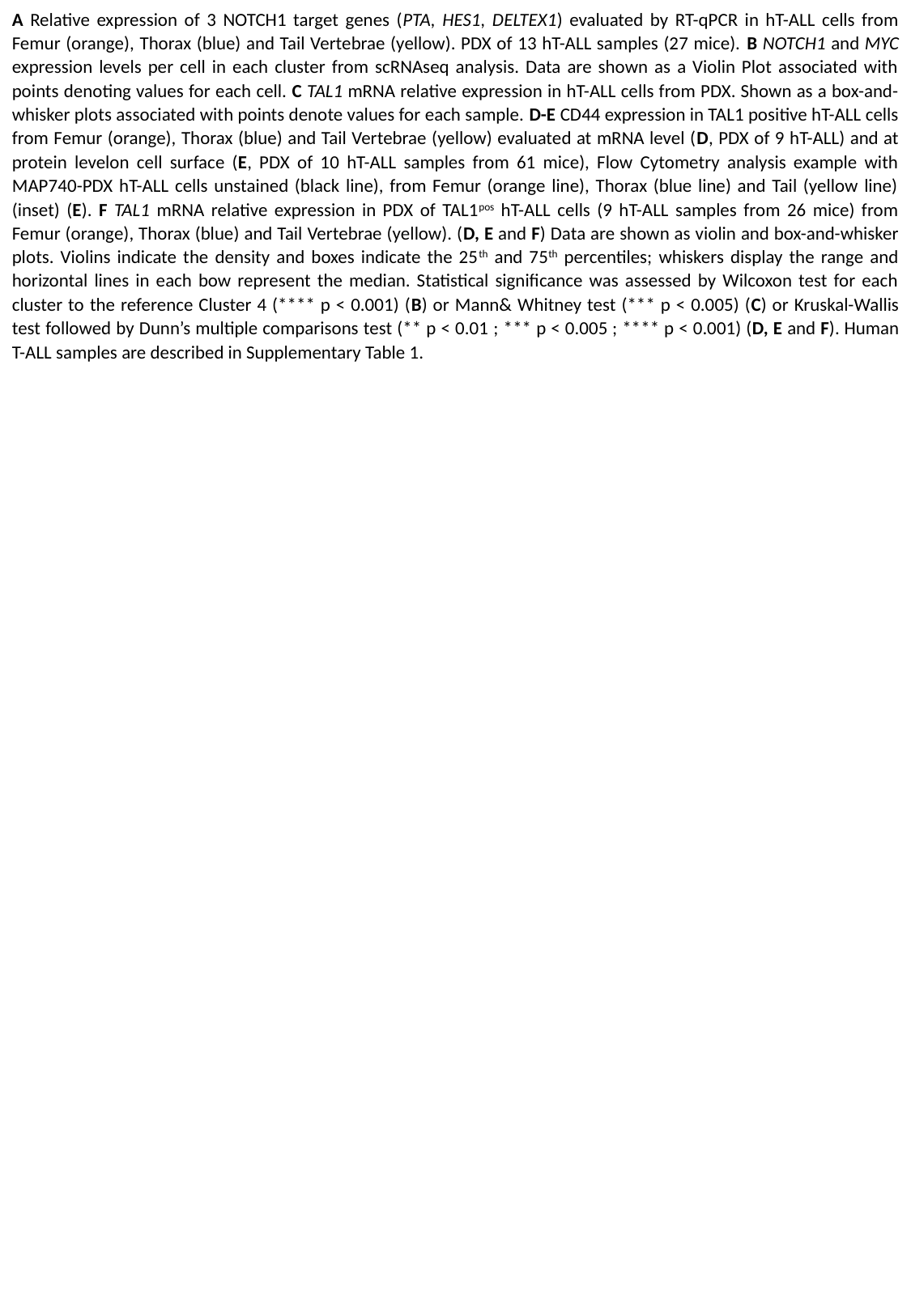

A Relative expression of 3 NOTCH1 target genes (PTA, HES1, DELTEX1) evaluated by RT-qPCR in hT-ALL cells from Femur (orange), Thorax (blue) and Tail Vertebrae (yellow). PDX of 13 hT-ALL samples (27 mice). B NOTCH1 and MYC expression levels per cell in each cluster from scRNAseq analysis. Data are shown as a Violin Plot associated with points denoting values for each cell. C TAL1 mRNA relative expression in hT-ALL cells from PDX. Shown as a box-and-whisker plots associated with points denote values for each sample. D-E CD44 expression in TAL1 positive hT-ALL cells from Femur (orange), Thorax (blue) and Tail Vertebrae (yellow) evaluated at mRNA level (D, PDX of 9 hT-ALL) and at protein levelon cell surface (E, PDX of 10 hT-ALL samples from 61 mice), Flow Cytometry analysis example with MAP740-PDX hT-ALL cells unstained (black line), from Femur (orange line), Thorax (blue line) and Tail (yellow line) (inset) (E). F TAL1 mRNA relative expression in PDX of TAL1^pos^ hT-ALL cells (9 hT-ALL samples from 26 mice) from Femur (orange), Thorax (blue) and Tail Vertebrae (yellow). (D, E and F) Data are shown as violin and box-and-whisker plots. Violins indicate the density and boxes indicate the 25^th^ and 75^th^ percentiles; whiskers display the range and horizontal lines in each bow represent the median. Statistical significance was assessed by Wilcoxon test for each cluster to the reference Cluster 4 (**** p < 0.001) (B) or Mann& Whitney test (*** p < 0.005) (C) or Kruskal-Wallis test followed by Dunn’s multiple comparisons test (** p < 0.01 ; *** p < 0.005 ; **** p < 0.001) (D, E and F). Human T-ALL samples are described in Supplementary Table 1.

### Slide 9
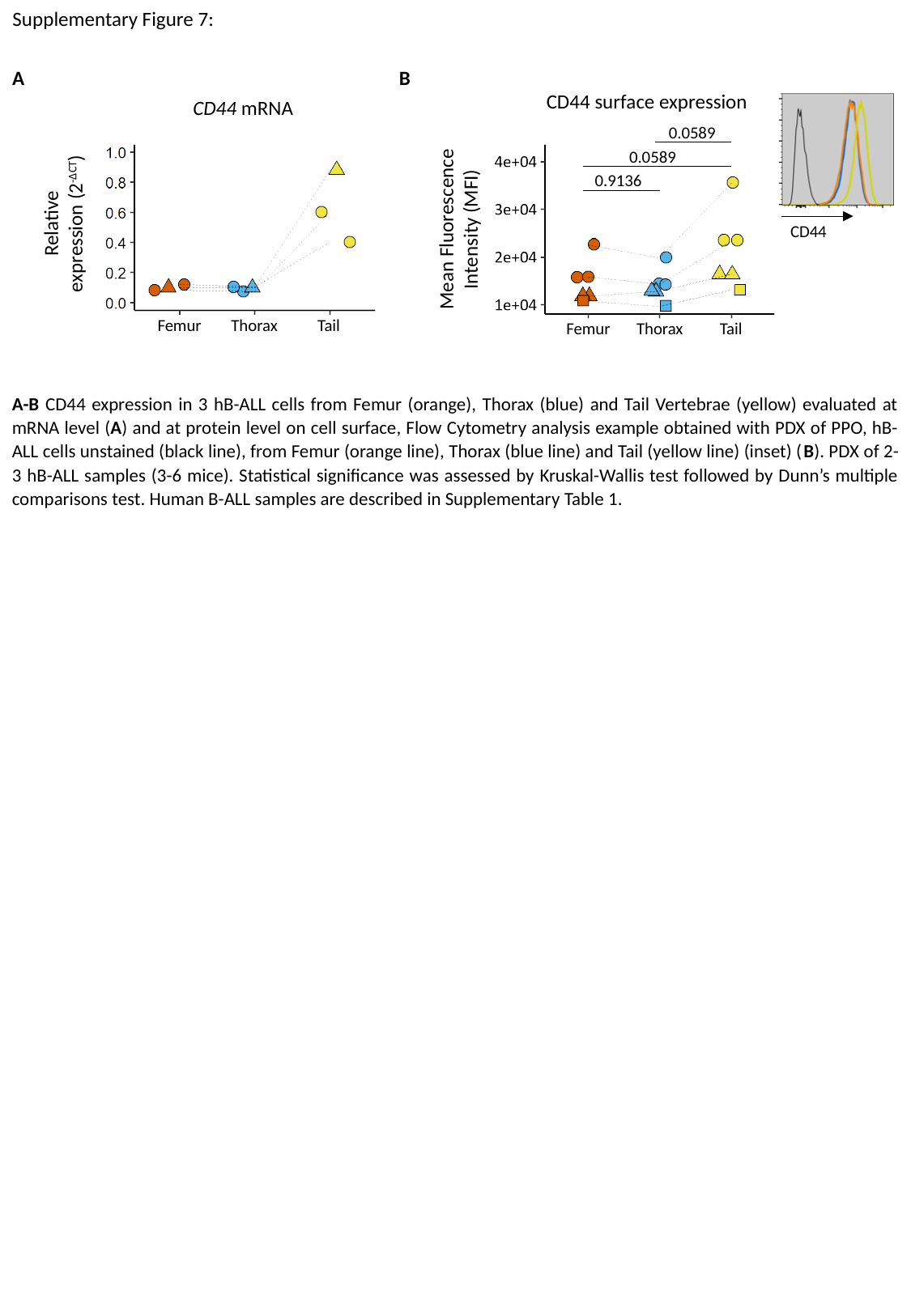

Supplementary Figure 7:
A
B
CD44 surface expression
0.0589
0.0589
0.9136
Mean Fluorescence
Intensity (MFI)
Femur
Thorax
Tail
CD44
CD44 mRNA
Relative expression (2^-ΔCT^)
Femur
Thorax
Tail
A-B CD44 expression in 3 hB-ALL cells from Femur (orange), Thorax (blue) and Tail Vertebrae (yellow) evaluated at mRNA level (A) and at protein level on cell surface, Flow Cytometry analysis example obtained with PDX of PPO, hB-ALL cells unstained (black line), from Femur (orange line), Thorax (blue line) and Tail (yellow line) (inset) (B). PDX of 2-3 hB-ALL samples (3-6 mice). Statistical significance was assessed by Kruskal-Wallis test followed by Dunn’s multiple comparisons test. Human B-ALL samples are described in Supplementary Table 1.

### Slide 10
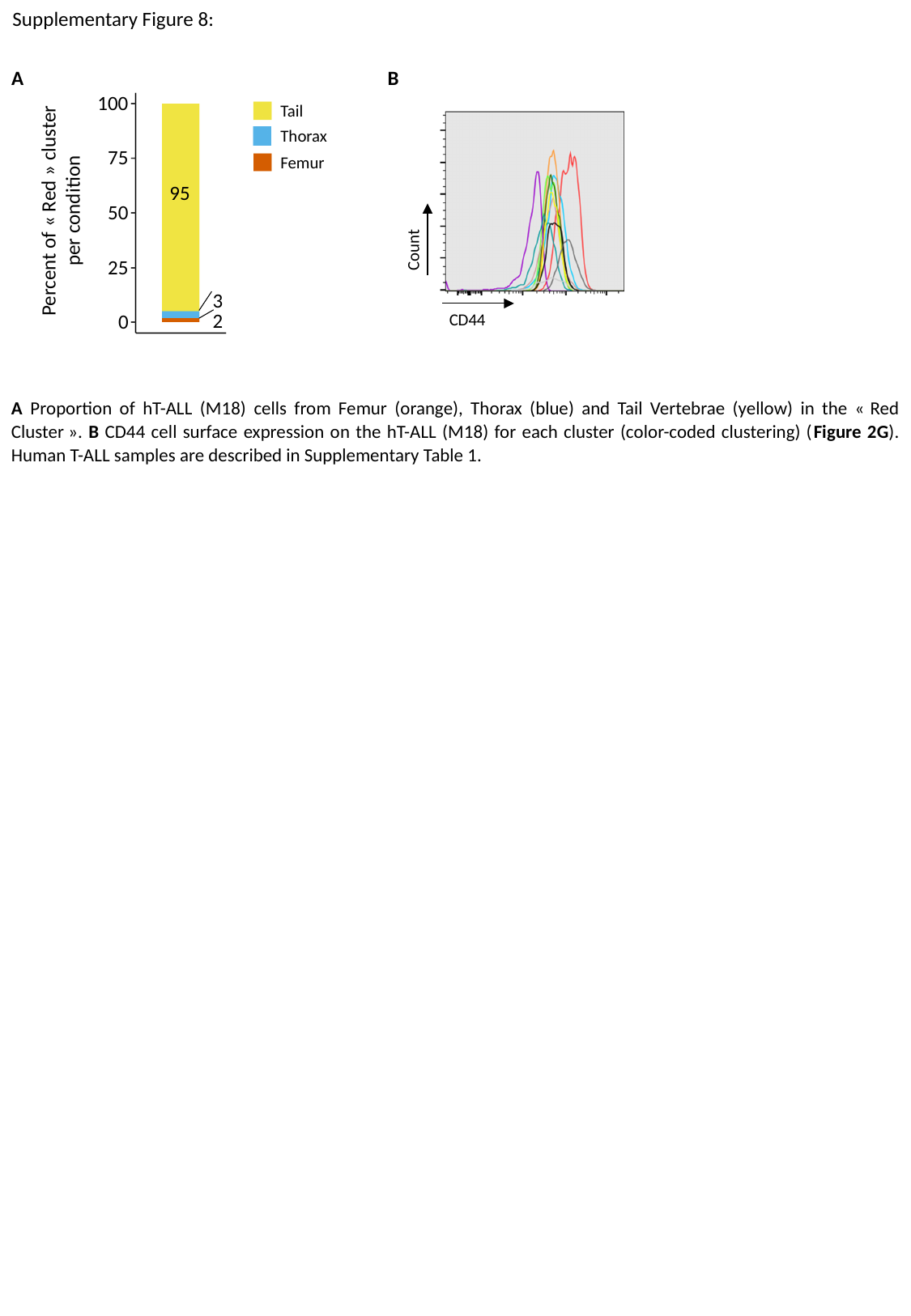

Supplementary Figure 8:
A
B
100
75
50
25
0
Tail
Thorax
Femur
95
Percent of « Red » cluster per condition
3
2
Count
CD44
A Proportion of hT-ALL (M18) cells from Femur (orange), Thorax (blue) and Tail Vertebrae (yellow) in the « Red Cluster ». B CD44 cell surface expression on the hT-ALL (M18) for each cluster (color-coded clustering) (Figure 2G). Human T-ALL samples are described in Supplementary Table 1.

### Slide 11
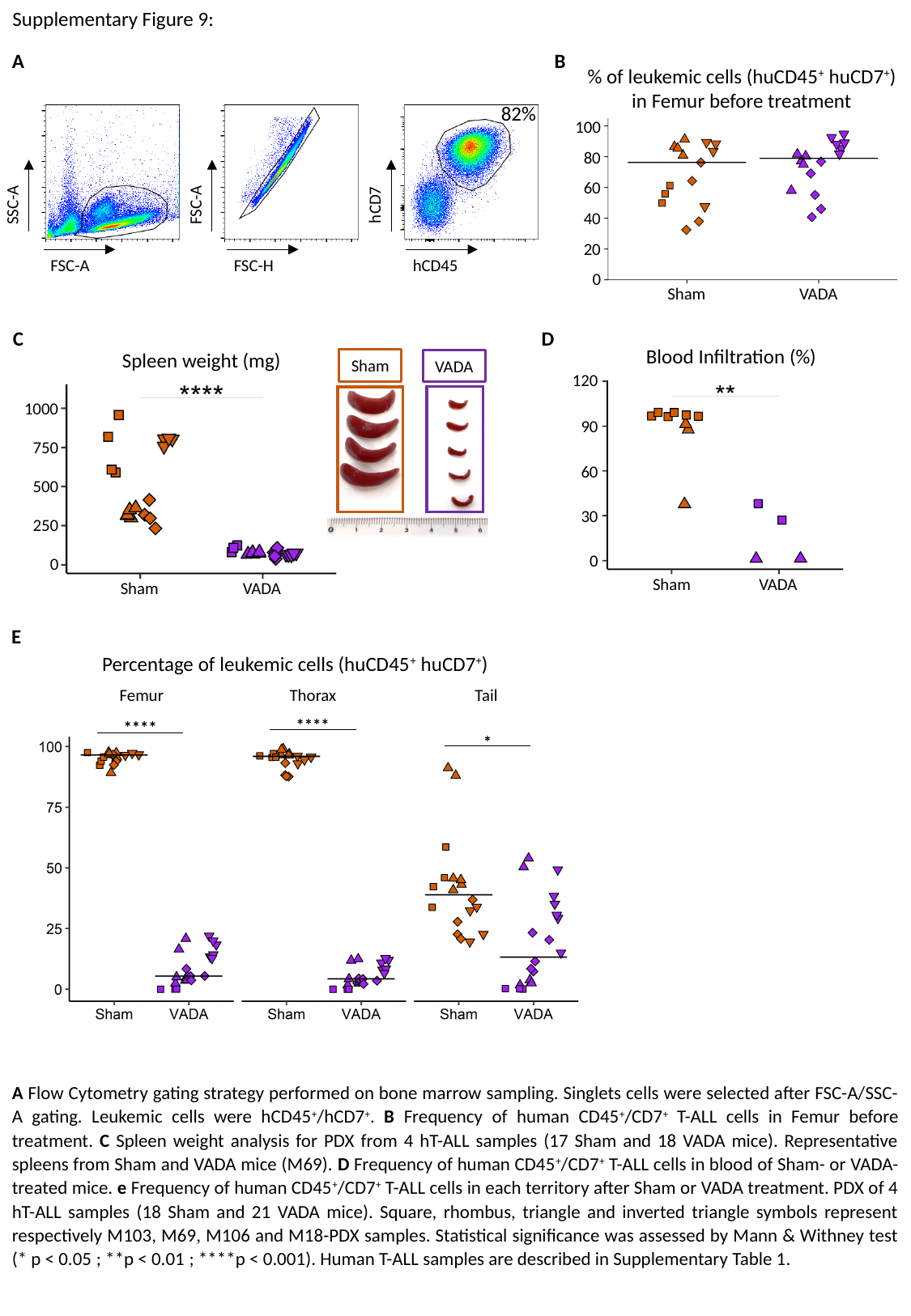

Supplementary Figure 9:
A
B
% of leukemic cells (huCD45^+^ huCD7^+^) in Femur before treatment
100
80
60
40
20
0
Sham
VADA
82%
SSC-A
FSC-A
FSC-A
FSC-H
hCD7
hCD45
C
Spleen weight (mg)
Sham
VADA
1000
750
500
250
0
Sham
VADA
D
Blood Infiltration (%)
120
90
60
30
0
Sham
VADA
E
Percentage of leukemic cells (huCD45^+^ huCD7^+^)
Femur
Thorax
Tail
****
****
*
A Flow Cytometry gating strategy performed on bone marrow sampling. Singlets cells were selected after FSC-A/SSC-A gating. Leukemic cells were hCD45^+^/hCD7^+^. B Frequency of human CD45^+^/CD7^+^ T-ALL cells in Femur before treatment. C Spleen weight analysis for PDX from 4 hT-ALL samples (17 Sham and 18 VADA mice). Representative spleens from Sham and VADA mice (M69). D Frequency of human CD45^+^/CD7^+^ T-ALL cells in blood of Sham- or VADA-treated mice. e Frequency of human CD45^+^/CD7^+^ T-ALL cells in each territory after Sham or VADA treatment. PDX of 4 hT-ALL samples (18 Sham and 21 VADA mice). Square, rhombus, triangle and inverted triangle symbols represent respectively M103, M69, M106 and M18-PDX samples. Statistical significance was assessed by Mann & Withney test (* p < 0.05 ; **p < 0.01 ; ****p < 0.001). Human T-ALL samples are described in Supplementary Table 1.

### Slide 12
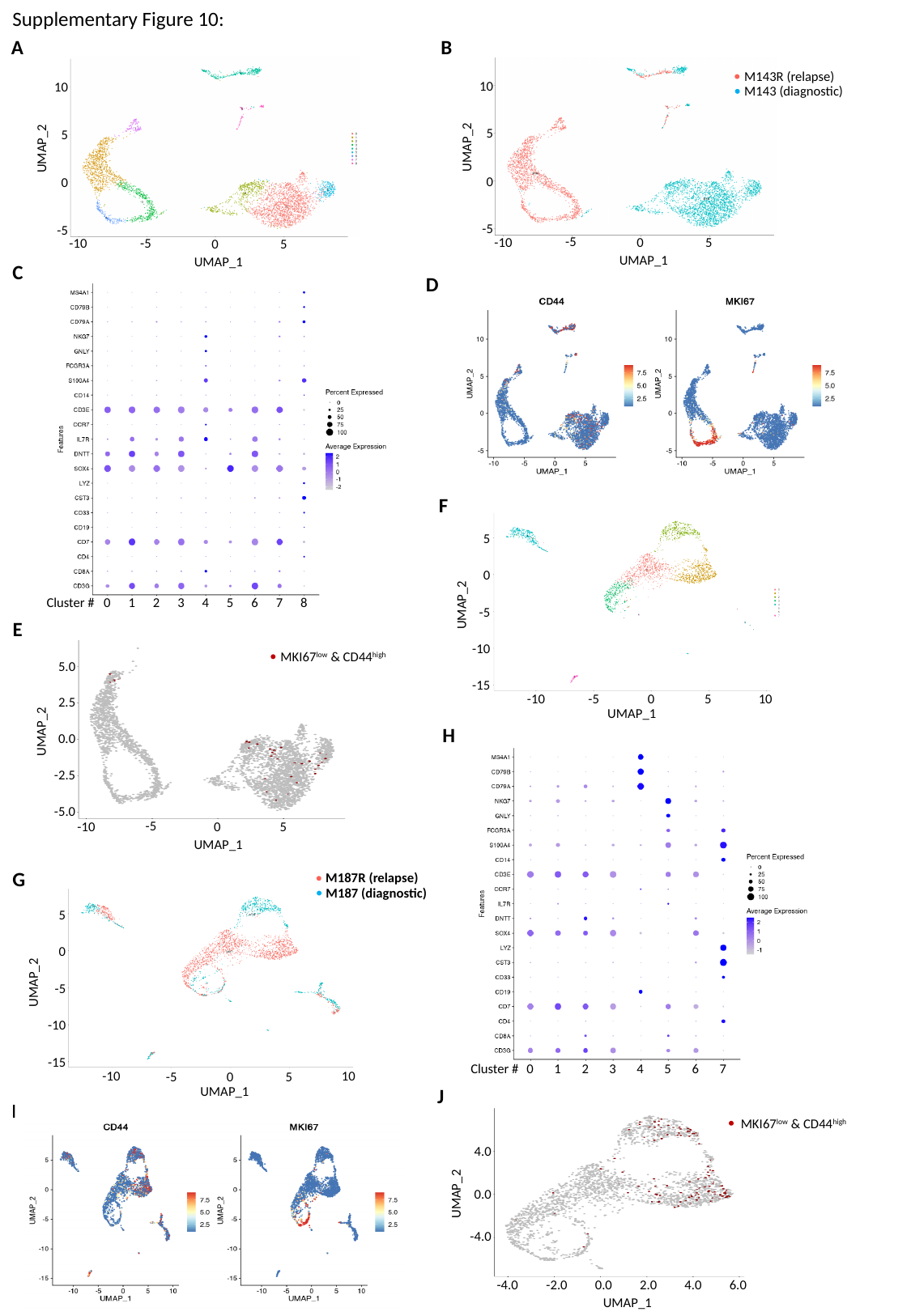

Supplementary Figure 10:
A
B
10
5
UMAP_2
0
-5
-5
-10
0
5
UMAP_1
M143R (relapse)
M143 (diagnostic)
10
5
UMAP_2
0
-5
-5
-10
0
5
UMAP_1
C
Cluster #
0
1
2
3
4
5
6
7
8
D
F
5
0
UMAP_2
-5
-10
-15
-10
-5
0
5
10
UMAP_1
E
5.0
2.5
UMAP_2
0.0
-2.5
-5.0
-5
-10
0
5
UMAP_1
MKI67^low^ & CD44^high^
H
Cluster #
0
1
2
3
4
5
6
7
G
M187R (relapse)
M187 (diagnostic)
5
0
UMAP_2
-5
-10
-15
-10
10
-5
0
5
UMAP_1
J
4.0
0.0
UMAP_2
-4.0
-4.0
0.0
4.0
UMAP_1
-2.0
2.0
6.0
MKI67^low^ & CD44^high^
I

### Slide 13
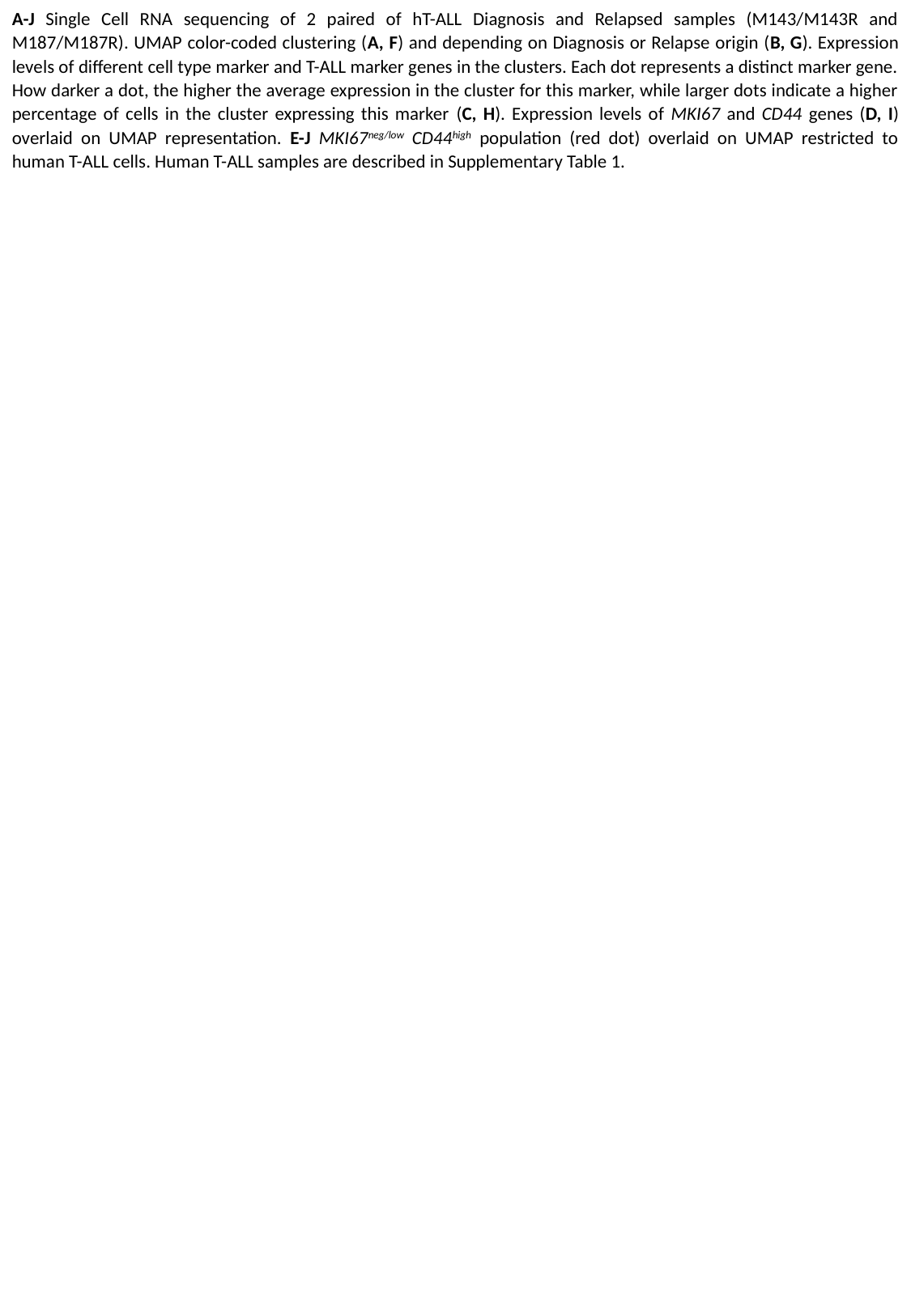

A-J Single Cell RNA sequencing of 2 paired of hT-ALL Diagnosis and Relapsed samples (M143/M143R and M187/M187R). UMAP color-coded clustering (A, F) and depending on Diagnosis or Relapse origin (B, G). Expression levels of different cell type marker and T-ALL marker genes in the clusters. Each dot represents a distinct marker gene. How darker a dot, the higher the average expression in the cluster for this marker, while larger dots indicate a higher percentage of cells in the cluster expressing this marker (C, H). Expression levels of MKI67 and CD44 genes (D, I) overlaid on UMAP representation. E-J MKI67^neg/low^ CD44^high^ population (red dot) overlaid on UMAP restricted to human T-ALL cells. Human T-ALL samples are described in Supplementary Table 1.

### Slide 14
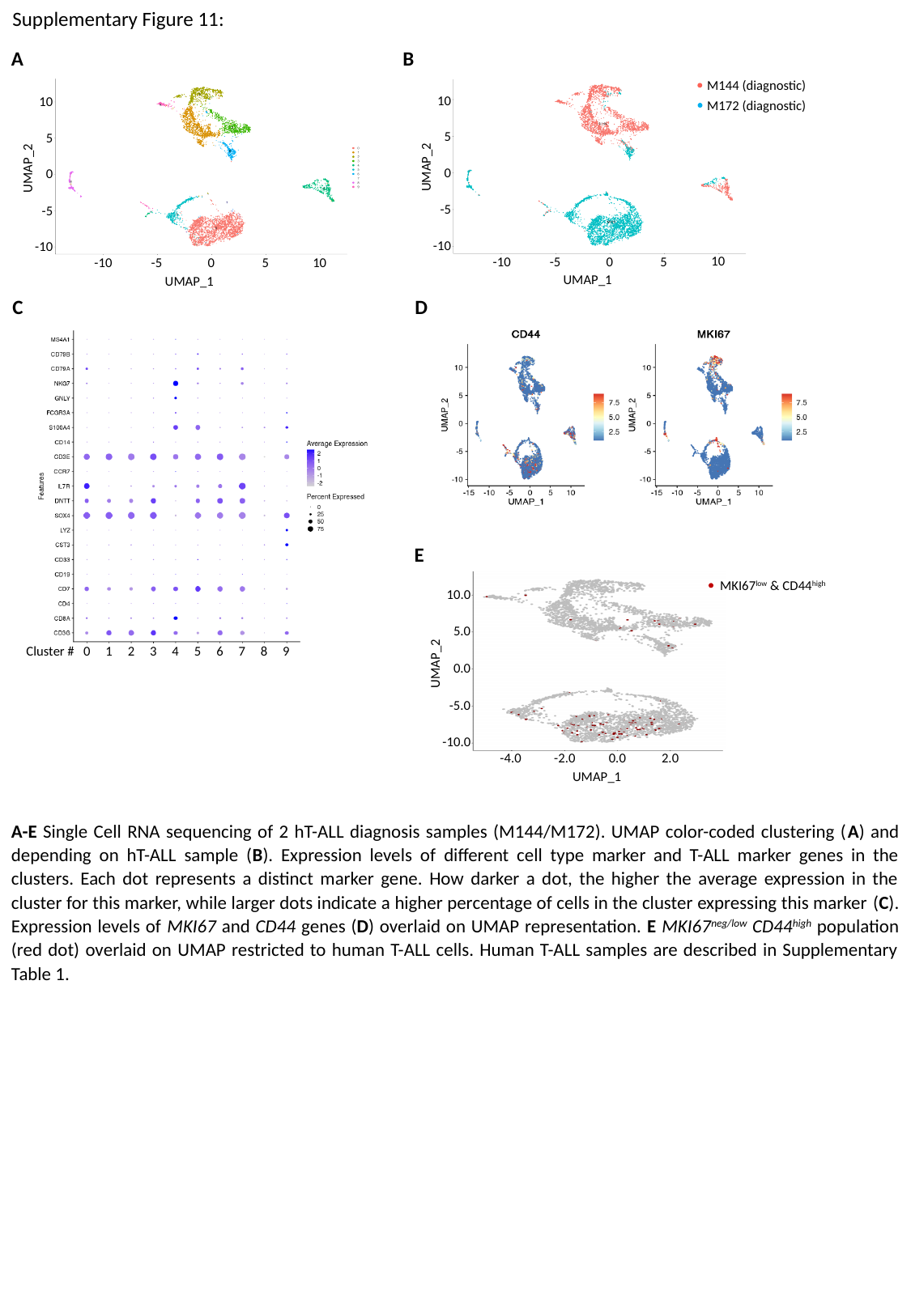

Supplementary Figure 11:
A
B
M144 (diagnostic)
M172 (diagnostic)
10
5
UMAP_2
0
-5
-10
-10
-5
0
5
UMAP_1
10
10
5
UMAP_2
0
-5
-10
-10
-5
0
5
UMAP_1
10
C
D
Cluster #
0
1
2
3
4
5
6
7
8
9
E
10.0
5.0
UMAP_2
0.0
-5.0
-10.0
-4.0
-2.0
0.0
2.0
UMAP_1
MKI67^low^ & CD44^high^
A-E Single Cell RNA sequencing of 2 hT-ALL diagnosis samples (M144/M172). UMAP color-coded clustering (A) and depending on hT-ALL sample (B). Expression levels of different cell type marker and T-ALL marker genes in the clusters. Each dot represents a distinct marker gene. How darker a dot, the higher the average expression in the cluster for this marker, while larger dots indicate a higher percentage of cells in the cluster expressing this marker (C). Expression levels of MKI67 and CD44 genes (D) overlaid on UMAP representation. E MKI67^neg/low^ CD44^high^ population (red dot) overlaid on UMAP restricted to human T-ALL cells. Human T-ALL samples are described in Supplementary Table 1.

### Slide 15
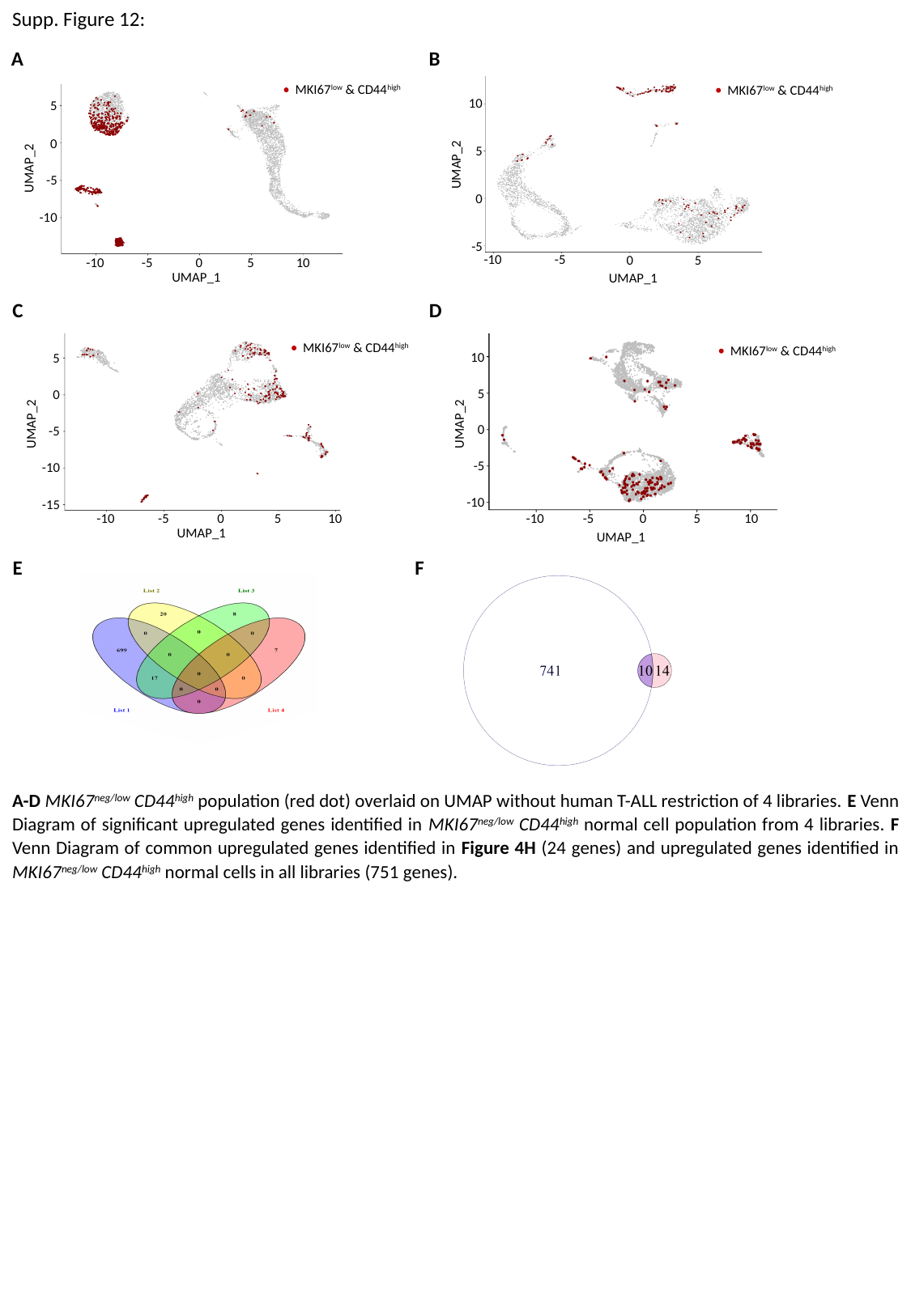

Supp. Figure 12:
A
B
MKI67^low^ & CD44^high^
5
0
UMAP_2
-5
-10
-10
-5
0
5
10
UMAP_1
MKI67^low^ & CD44^high^
10
5
UMAP_2
0
-5
-5
-10
0
5
UMAP_1
C
D
MKI67^low^ & CD44^high^
10
5
UMAP_2
0
-5
-10
-10
-5
0
5
UMAP_1
10
MKI67^low^ & CD44^high^
5
0
UMAP_2
-5
-10
-15
-10
-5
0
5
10
UMAP_1
E
F
A-D MKI67^neg/low^ CD44^high^ population (red dot) overlaid on UMAP without human T-ALL restriction of 4 libraries. E Venn Diagram of significant upregulated genes identified in MKI67^neg/low^ CD44^high^ normal cell population from 4 libraries. F Venn Diagram of common upregulated genes identified in Figure 4H (24 genes) and upregulated genes identified in MKI67^neg/low^ CD44^high^ normal cells in all libraries (751 genes).

### Slide 16
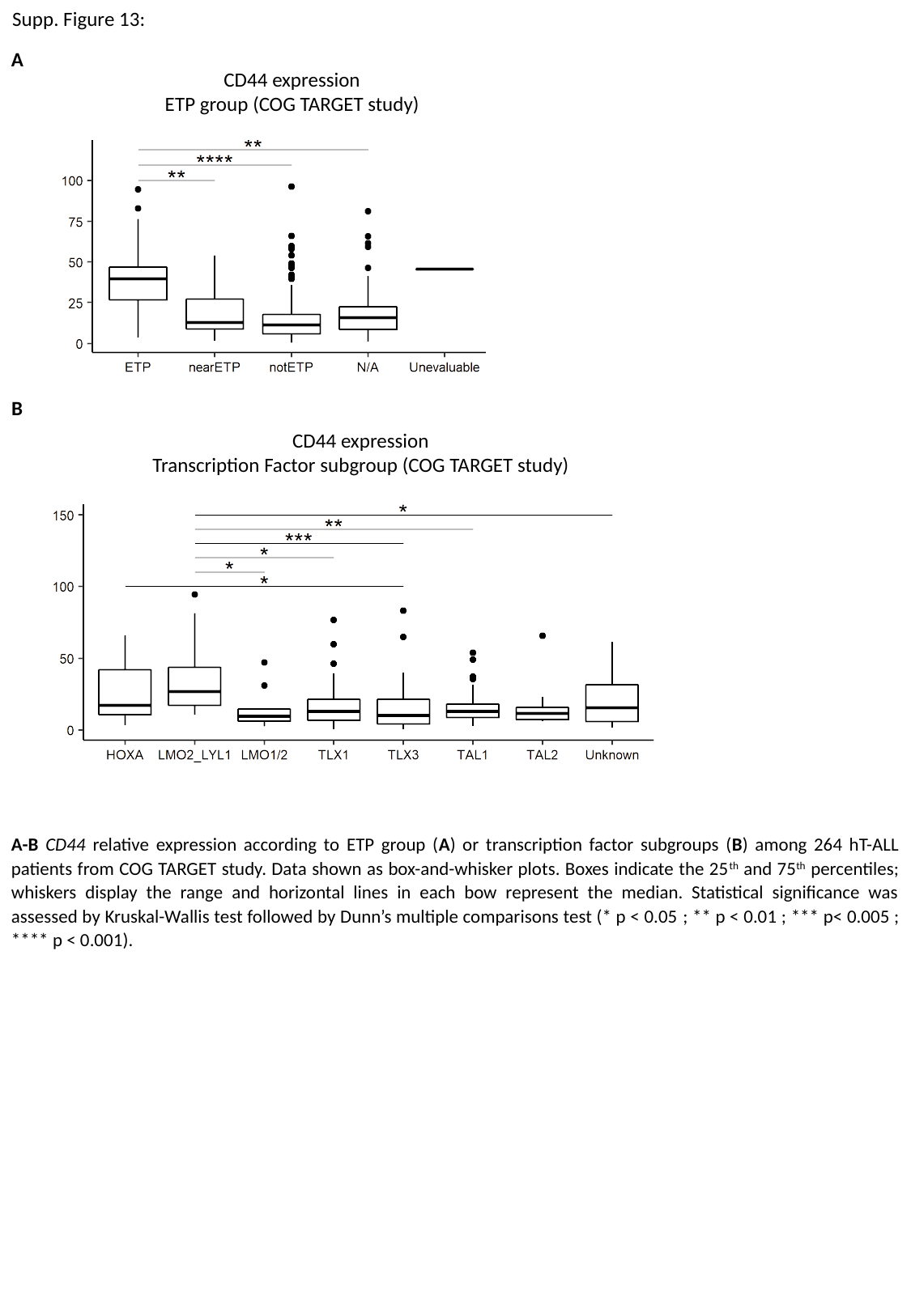

Supp. Figure 13:
A
CD44 expression
ETP group (COG TARGET study)
B
CD44 expression
Transcription Factor subgroup (COG TARGET study)
A-B CD44 relative expression according to ETP group (A) or transcription factor subgroups (B) among 264 hT-ALL patients from COG TARGET study. Data shown as box-and-whisker plots. Boxes indicate the 25^th^ and 75^th^ percentiles; whiskers display the range and horizontal lines in each bow represent the median. Statistical significance was assessed by Kruskal-Wallis test followed by Dunn’s multiple comparisons test (* p < 0.05 ; ** p < 0.01 ; *** p< 0.005 ; **** p < 0.001).

### Slide 17
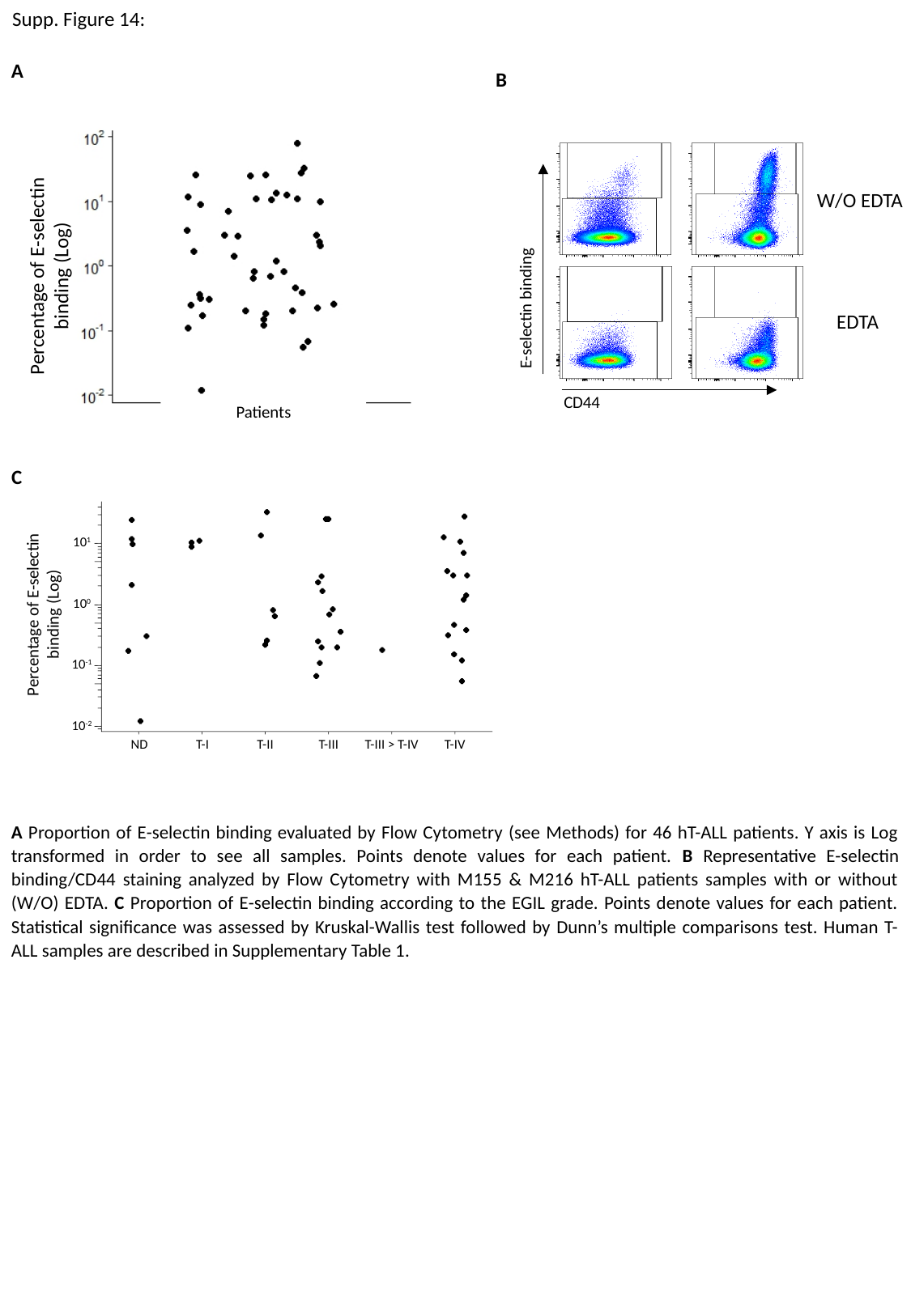

Supp. Figure 14:
A
B
Percentage of E-selectin binding (Log)
Patients
W/O EDTA
E-selectin binding
EDTA
CD44
C
10^1^
Percentage of E-selectin binding (Log)
10^0^
10^-1^
10^-2^
ND
T-I
T-II
T-III
T-III > T-IV
T-IV
A Proportion of E-selectin binding evaluated by Flow Cytometry (see Methods) for 46 hT-ALL patients. Y axis is Log transformed in order to see all samples. Points denote values for each patient. B Representative E-selectin binding/CD44 staining analyzed by Flow Cytometry with M155 & M216 hT-ALL patients samples with or without (W/O) EDTA. C Proportion of E-selectin binding according to the EGIL grade. Points denote values for each patient. Statistical significance was assessed by Kruskal-Wallis test followed by Dunn’s multiple comparisons test. Human T-ALL samples are described in Supplementary Table 1.
